# Understanding the binding mechanisms of Alzheimer risk factor TREM2 in lipid bilayer using multi-scale molecular dynamics simulations

**DOI:** 10.1101/2024.03.08.584168

**Authors:** Zhiwen Zhong, Martin Ulmschneider, Christian D. Lorenz

## Abstract

Alzheimer’s disease (AD) is a widespread neurodegenerative condition affecting millions globally. Recent research has implicated variants of the triggering receptor expressed in myeloid cells 2 (TREM2) as risk factors for AD. TREM2, an immunomodulatory receptor on microglial surfaces, plays a pivotal role in regulating microglial activation by associating with DNAX-activation protein 12 (DAP12). Despite its significance, the mechanism underlying the formation of the complex between the transmembrane domains (TMDs) of TREM2 and DAP12 remains unclear. This study employs multi-scale molecular dynamics (MD) simulations to investigate three TMD complex models, including two derived from experiments and one generated by AlphaFold2. Conducted within a lipid membrane consisting of an 80:20 mixture of phosphatidylcholine (POPC) and cholesterol, our analysis reveals hydrogen bonding interactions between K26 of TREM2 and D16 of DAP12 in all three models, consistent with previous experimental findings. Our results elucidate the different spatial conformations observed in the models and offer insights into the structure of the TREM2/DAP12 TMD complex. Furthermore, we elucidate the role of charged residues in the assembly structure of the complex within the lipid membrane. These findings enhance our understanding of the molecular mechanism governing TREM2/DAP12 complex formation, providing a foundation for designing novel therapeutic strategies to address AD and other neu-rodegenerative diseases.

**Graphical Abstract:** 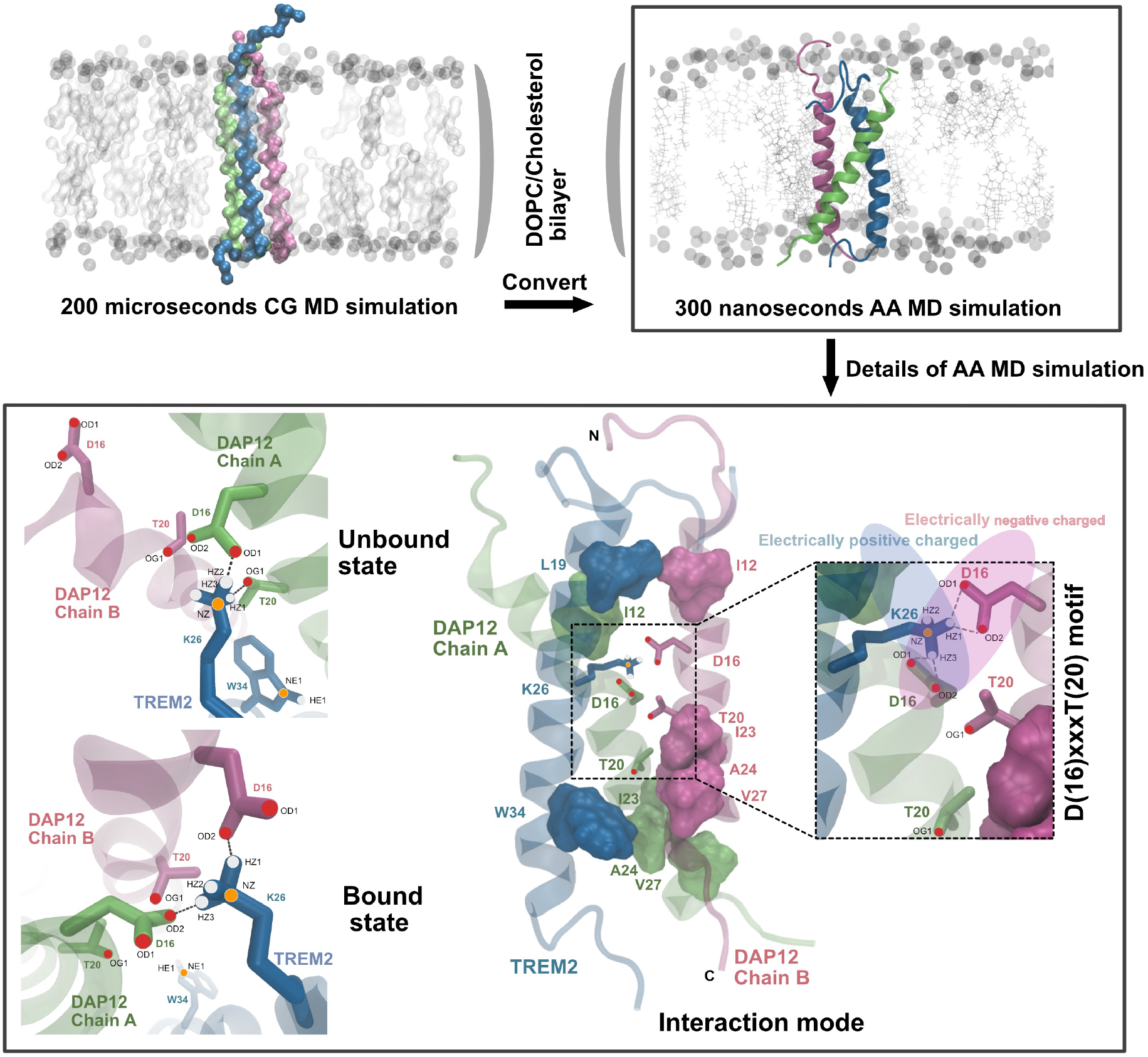

**Highlights:**

- Different TREM2/DAP12 complexes are generated using experimental PDB structures or AlphaFold2
- Long time coarse-grained MD simulations are used for getting the systems stable
- Atomistic detailed complex structures are captured using all-atom sim-ulation
- Provides mechanistic insight into TREM2 transformation from its unbound state to bounded state
- Offers novel insights into the molecular basis of TREM2/DAP12 sig-nalling pathway

## Introduction

Alzheimer’s disease (AD) is a prevalent neurodegenerative disease that is the leading cause of dementia in the elderly. ^1^ It was first identified over a century ago by the German neuropsychiatrist, Alois Alzheimer,^2^ and now has been proven to have a complicated genetic architecture.^3^ The encoding genes, such as presenilin-1 (PSEN1) and presenilin-2 (PSEN2), which are part of the *γ*-secretase complex, along with the *ɛ*4 allele of apolipoprotein E (APOE) and amyloid precursor protein (APP), exhibit a high degree of mutation penetrance. Amyloid plaques are one of the hallmarks of AD ^4^ and are mainly compromised of *β*-amyloid peptides (A*β*s) which are derived from the APP.^5^ The Triggering Receptor Expressed on Myeloid Cells 2 (TREM2) is a transmembrane protein that is thought to be related to AD in two significant ways. First, whole-genome analyses have identified rare variants in TREM2 that increase the risk of AD by approximately threefold,^6,7^ similar to APOE. Secondly, TREM2 is selectively expressed in myeloid cells, where it promotes the optimal microglial function required to attenuate AD progression. Microglia, which are the primary component of the human brain’s immune system, are involved in A*β* accumulation and thus contribute to the development of AD.^8^ Thus, TREM2 is a potential target for eliciting a protective role for microglia in AD and other neurodegenerative diseases. DAP12 (DNAX-activating protein of 12 kDa) is a transmembrane protein that activates downstream signalling pathways by binding to TREM2.^9,10^ Together, TREM2 and DAP12 play crucial roles in the regulation of microglia function, including phagocytosis and inflammatory responses.

The initiation of the TREM2 signalling pathway relies on a stable interaction between TREM2 and DAP12 within the membrane. Researchers proposed that TREM2 could bind to numerous molecules, and its signalling intensity and direction could be differentially influenced by interacting with various ligands.^11^ The TREM2 pathway is first triggered when the N-terminus of TREM2 binds to ligands in the extracellular domain, which activates downstream pathways that involve a stable interaction between TREM2 and DAP10/12.^12^ Then the ectodomain is cleaved off by metalloproteinase domain-containing proteins 10/17 (ADAM 10/17), and the C-terminal fragment (CTF) is cleaved by *γ*-secretase (a *γ*-secretase high-resolution cryo-EM structure has recently been solved^13^), ultimately terminating the TREM2 pathway.^14,15^ The termination of the TREM2 signalling pathway allows microglia to return to their basal state. Mutations in TREM2 and DAP12 genes have been linked to an increased risk of developing neurodegenerative diseases such as Alzheimer’s disease.^16^ Understanding the molecular mechanisms underlying the TREM2 signalling pathway and associated signalling adaptor molecules such as DAP12 is critical for the development of new therapeutic strategies for neurodegenerative diseases.^17,18^ Further research on this pathway will provide valuable insights into the regulation of microglia function and the potential development of new drugs targeting neurodegenerative diseases.

Recent studies have revealed that TREM2 binds to the DAP12 dimer via salt bridges that involve an aspartic acid on each chain of the DAP12 protein and a lysine in TREM2, and therefore ultimately forms a trimer.^19^ The orientation of the transmembrane domain of the TREM2 monomer and the DAP12 dimer extends from the N-terminus to the C-terminus, spanning from the outer cell membrane to the inner cell membrane. Furthermore, researchers have demonstrated that the TREM2 transmembrane helix is stabilized by either binding to DAP12 or as a result of the mutation of the charged amino acid K26 to an alanine (K26A).^20^ The presence of a charged amino acid (K26) in the transmembrane domain leads to the adoption of a kinked structure by the TREM2 transmembrane domain (TMD), which exhibits greater flexibility. Conversely, the removal of this charge (K26A) results in the stabilization of the TREM2-TMD, thereby reducing its dynamics, and resulting in its structure being similar to when it is complexed with DAP12.^20^ While there is substantial interest in understanding how TREM2 recognizes DAP12 in the membrane, a comprehensive understanding of the atomistic interactions governing the formation of this stable complex is still lacking. Molecular dynamics (MD) simulations offer a distinct opportunity to foster this level of understanding. By modelling the inter-atomic interactions, MD simulations predict the movements of individual atoms in complex molecular systems, providing insights into fundamental biomolecular processes such as protein folding, ligand binding, and conformational changes.^21^ The growing accessibility of experimentally determined structures, exemplified by repositories like the Protein Data Bank (PDB),^22^ the evolution of machine learning techniques for predicting protein structures (such as AlphaFold2), ^23^ and the continuous enhancement of computational hardware have significantly bolstered the prominence of MD simulations. These advancements facilitate the simulation of larger and more intricate molecular systems within reduced timeframes.

In this study, we studied three models of the TREM2/DAP12 (TD) complex in a phosphatidylcholine (POPC) and cholesterol (80:20) lipid environment: two based on experimental data and one modelled using AlphaFold2, involving TREM2/DAP12 complexes simulated within a phosphatidylcholine (POPC) and cholesterol (80:20) lipid environment (Figure 1). Employing long-timescale coarse-grain (CG) MD simulations to stabilize the systems, we subsequently converted them to all-atom (AA) MD simulations. Our atomic-level analysis revealed unique protein-protein interfaces in the three systems, resulting from a combination of salt bridges, hydrogen bonds, and hydrophobic interactions. Notably, all three systems showcased a K26/D16 interaction involving both a salt bridge and hydrogen bonds. This interaction appears to be crucial in the recognition process of the TD complex.

**Figure 1:**
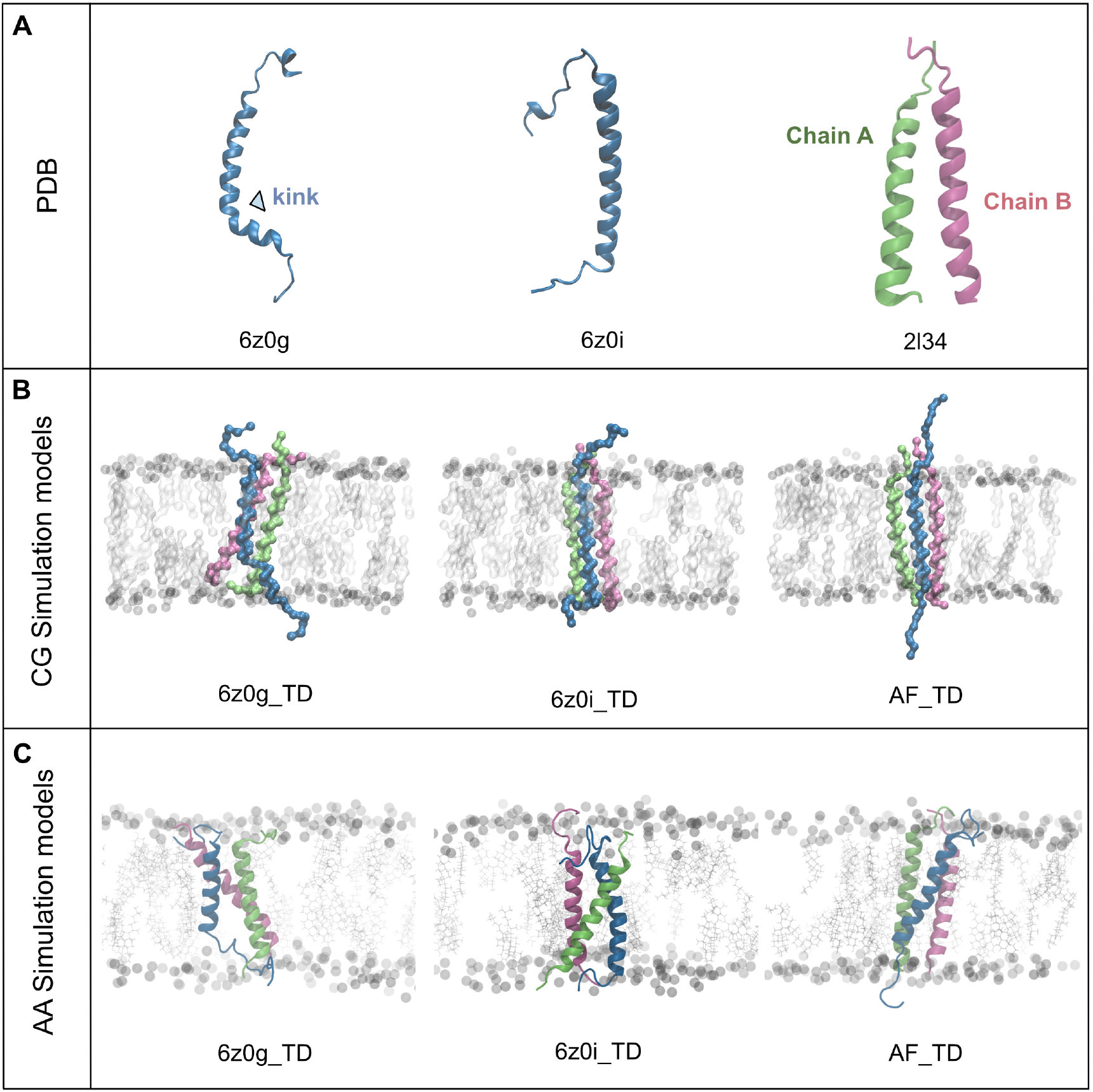
Representation of three TREM2/DAP12 (TD) transmembrane domain complexes in a lipid environment. (A) From left to right: 6z0g illustrates the kinked structure of TREM2, while 6z0i depicts the TREM2 PDB structure interacting with DAP12. The right-most structure portrays the DAP12 dimer, with Chain A in lime and Chain B in mauve. (B) Illustrates the coarse-grained 3D structure of the three models, where the grey sphere represents the POPC bilayer, and cholesterols are visualized using a licorice representation. Protein structure colours are consistent with (A). (C) Presents the all-atom 3D structures of the three models within the POPC/cholesterol environment. The grey sphere indicates the POPC bilayer, and silver dots denote cholesterol.

## Results

### Three TREM2/DAP12 Complexes in a Lipid Environment

We conducted a 200 *µ*s CG simulation and then converted the final configuration to an AA representation. Afterwards, we ran the AA system for an additional 300 ns within the same lipid environment. The system’s stability after the 200 *µ*s simulations was confirmed by assessing the total number of contacts between the proteins (Figure S1) and the contact maps between individual residues within the various proteins (Figure S2 – S4). The contact maps depicted consistent contact between residues in the central region of the different proteins across four different time frames of the trajectories.

Of the three systems we studied (Figure 1C), 6z0g represents the NMR structure of apo TREM2, while 6z0i depicts the NMR structure of TREM2 when it is bound with DAP12 (Figure 1A), although the TREM2 helix part alone was resolved. A notable difference between 6z0g and 6z0i in the three-dimensional structure is the presence of a kinked region in the TREM2 transmembrane domain. Subsequently, the DAP12 dimer (PDB:2l34) was assembled separately with the TREM2 monomers (PDB: 6z0g and 6z0i) and formed 6z0g TD and 6z0i TD systems respectively (Figure 1C). The third system, AF TD, was generated using AlphaFold2 and shares the same sequence as 6z0g TD and 6z0i TD. The confidence level of the AF TD complex is perceptible and demonstrated in Figure S5.

### Kink Angles in Three Systems Reveal Different Binding Mechanisms

Within all three systems, TREM2 exists as a single *α*-helix transmembrane protein, while DAP12 is a dimer. Notably, the 6z0g TD system exhibits a kink, while 6z0i TD and AF TD show no kinks, consistent with the apo PDB structures (Figure 1A). We determined the kink angle using four C*_α_* atoms in the TREM2 *α*-helix (shown in Figure 2A). During the 300 ns simulation, the average kink angle for 6z0g TD measured 99.8 *±* 0.1°, whereas 6z0i TD and AF TD showed kink angles of 37.7 ± 0.1° and 37.4 ± 0.1°, respectively (Figure 2B). Upon analyzing the kink angle distributions, we observed that the most frequent kink angle for 6z0g TD is approximately 104° (Figure 2C). In contrast, 6z0i TD and AF TD displayed less pronounced kinks, with the most prevalent angles measuring 38° and 40°, respectively. These findings suggest that after our extended simulations (200 *µ*s CG simulation and 300 ns AA simulation), the kink persisted in the 6z0g TD model, while the other two models exhibited less pronounced kinks. This implies that different TREM2-DAP12 binding modes exist among the three systems.

**Figure 2:**
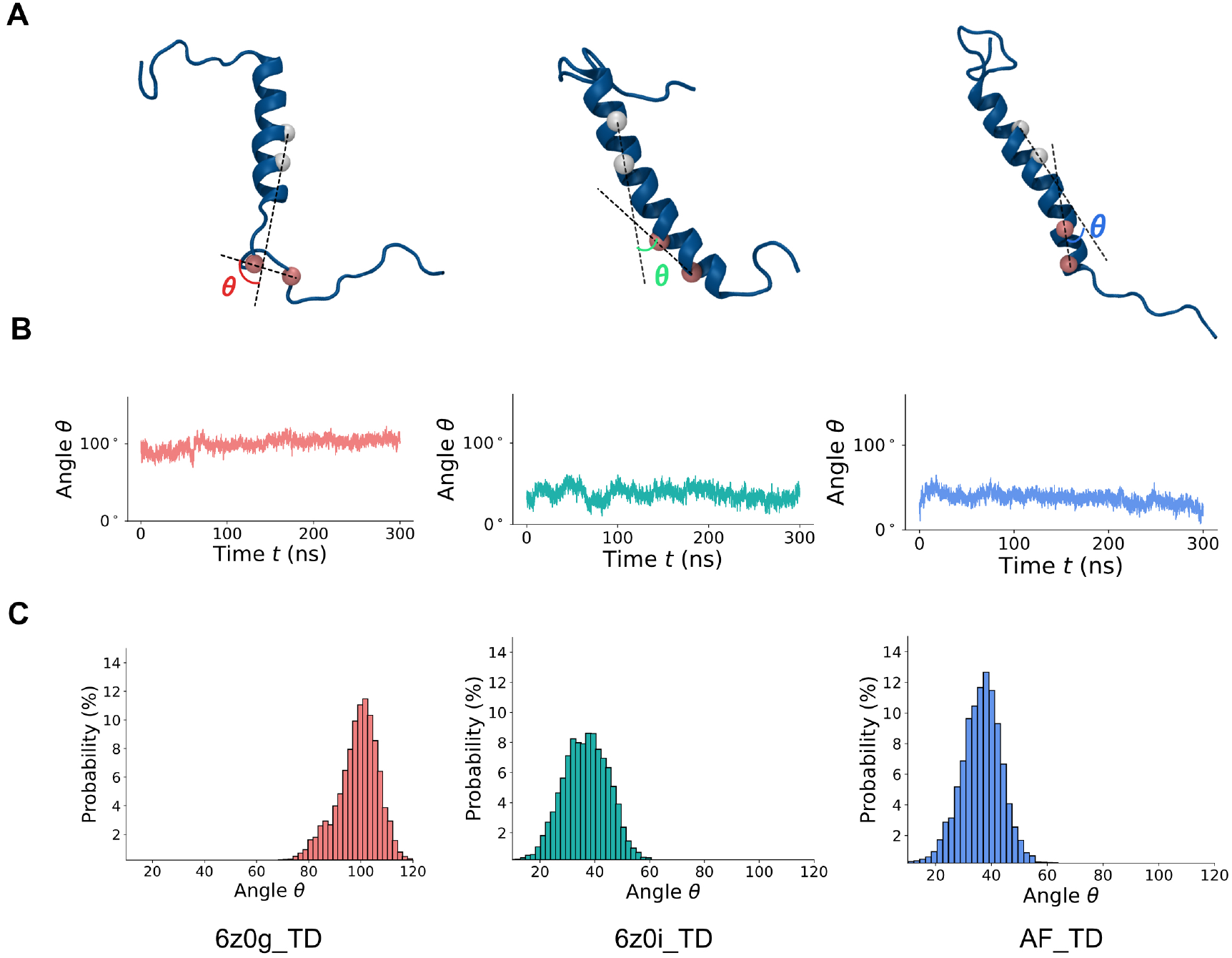
Representation of kink angles in three different systems. (A) Snapshots of TREM2 structures were taken at 150 ns. White spheres represent C*_α_* atoms of A20 and L24, while pink spheres represent C*_α_* atoms of A32 and A36. (B) Diagrams displaying the kink an-gles of the three models. Red, green, and blue lines correspond to 6z0g TD, 6z0i TD, and AF TD, respectively. (C) Illustrations showcasing the distribution of kink angles. Colors are consistent with (B).

### Contact Maps Using C***_α_*** Atoms Distance as Cut-off Reveal the Interaction between TREM2 and Two Chains of DAP12

We present representative snapshots and contact maps for TREM2 and both chains of the DAP12 dimer from each model (Figure 3). The kinked region around K26 in TREM2 and the DXXXT motif (where D and T represent D16 and T20 respectively, and X represents any amino acid) in DAP12 are known to have a salt bridge interaction. We observed that the middle region (the *α*-helix of the trimer, spanning residues 14 to 41 of TREM2 and 6 to 32 of DAP12, depicted by the light blue rectangle) of the unkinked models (6z0i TD & AF TD) of the TREM2 protein makes similar amounts of contact with the two DAP12 chains, while in the kinked model (6z0g TD), the middle region part of TREM2 only makes significant contact with one chain (Chain A). Therefore, despite the different models having identical amino acid sequences in Chain A and Chain B of DAP12, the kinked and unkinked models of TREM2 exhibit distinct binding preferences.

**Figure 3:**
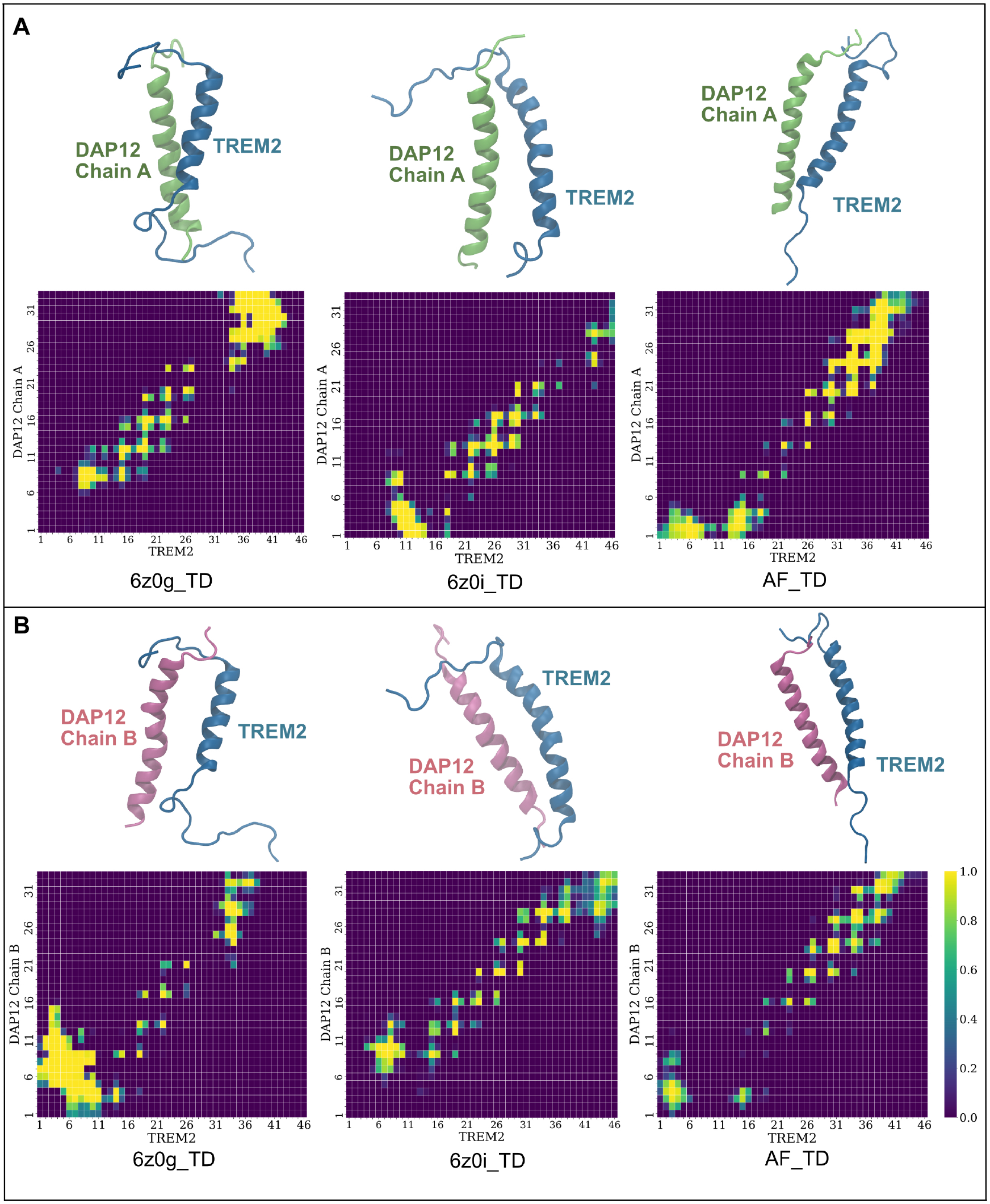
Snapshots of 3D structures and contact maps of three systems. The snapshot was taken at 200 ns, and the contact map was generated between 175 to 225 ns. (A) illustrates TREM2 contact with DAP12 Chain A in the three models. DAP12 Chain A structures are depicted in lime, while TREM2 is shown in blue. The contact map utilizes the “rocket” preset colour scheme. (B) showcases TREM2 contacts with DAP12 Chain B, with DAP12 Chain B represented in mauve and TREM2 in blue.

### Identification of Potential Binding Residues in three TREM2/DAP12 Complexes

To gain deeper insights into the various binding mechanisms, we constructed contact maps using a 3 Å cut-off distance, involving all atoms including side chain atoms (Figure S6). Contacts persisting for more than 50% of the total simulation time were considered in the normalized contact map (Figure S7). Summarizing the six contact maps, we illustrated all contacts in Figure 4A. From this data, the most common contacts between residues, which are those that were present in equal or more than half of the six contact maps, were plotted in Figure 4B.

**Figure 4:**
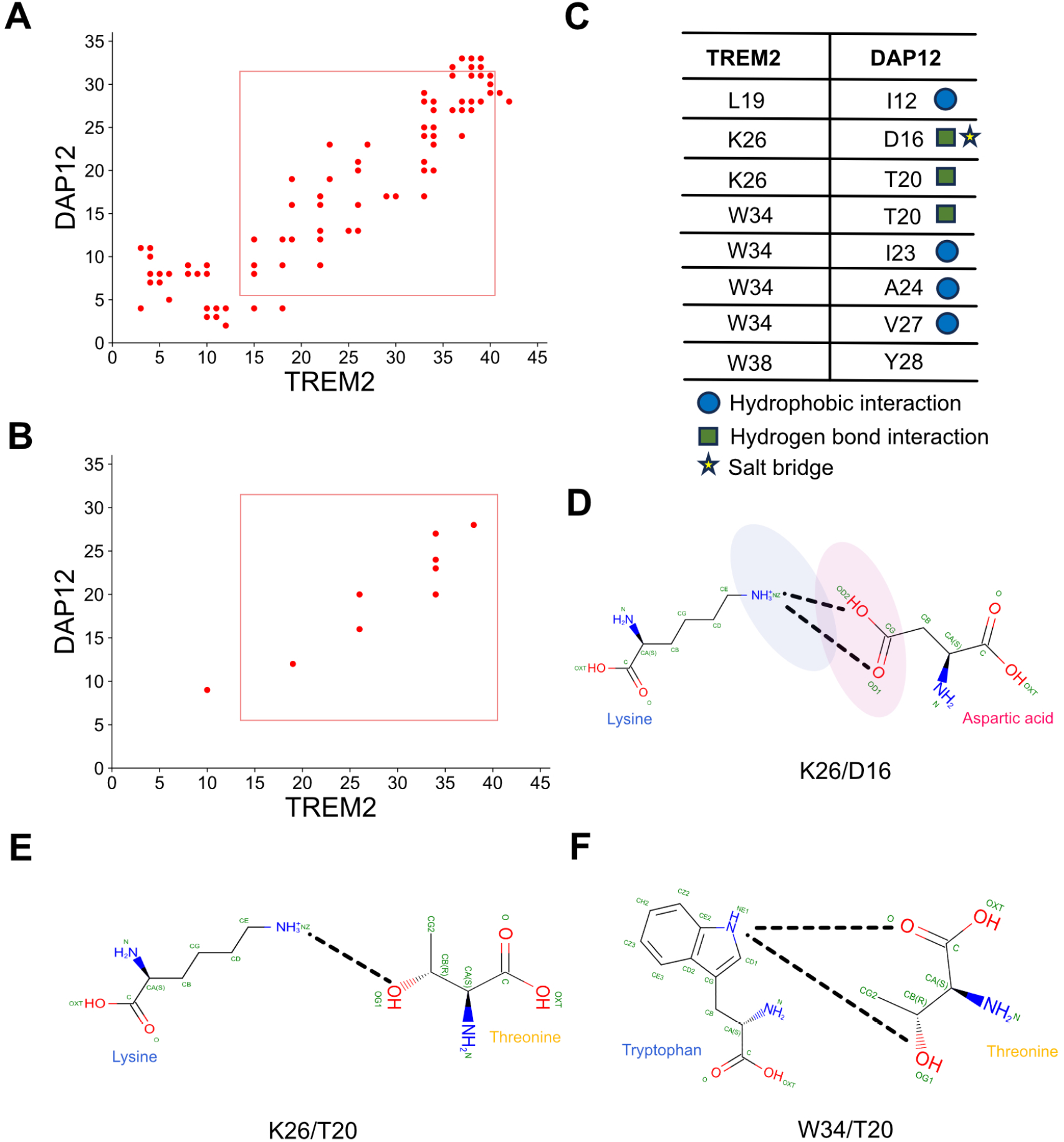
Representation of key binding residues between TREM2 and DAP12. (A) displays merged total contacts. The rectangle indicates the *α*-helix region of the TD complex. (B) presents the calculated contact map. (C) illustrates the key binding residue pairs of TREM2 and DAP12. The blue circle represents hydrophobic interactions, the green rectangle signifies hydrogen bond interactions, and the yellow star denotes salt bridge interactions. (D) demonstrates the binding of K26 of TREM2 to D16 of DAP12. The dotted black line represents hydrogen bonds between relative atoms. The blue oval region signifies the positively charged area, while the pink oval region indicates the negatively charged area. (E) showcases the binding of K26 to T20 and the hydrogen bonds between them. (F) exhibits the binding of W34 to T20 and the hydrogen bonds between them.

Upon analyzing these pivotal pair of residues that are in contact in Figure 4B, we inferred that hydrophobic interactions, hydrogen bonds, and salt bridge interactions play crucial roles in the binding of TREM2 to DAP12 (Figure 4C). Furthermore, we produced atomistic contact maps for each of the eight significant residue pairs (Figures S8 - S15). These atomistic contact maps help delineate the specific atoms that are in contact within the identified residues.

### Analysis of Hydrogen Bonds in the TREM2/DAP12 Interaction

Initially, we focussed on three potential hydrogen bond partners within the TREM2/DAP12 complex (K26/D16, K26/T20, and W34/T20; Figures 4D, E & F, respectively) and investigated the distances between the donors and acceptors involved in each of the hydrogen bonds (Figure 5). Analyzing the potential K26/D16 hydrogen bonds in 6z0g TD revealed that the distances between the NZ in K26 of TREM2 and OD1 and OD2 in D16 in Chain B of DAP12 (Figure 4D) are larger than 12 Å, exceeding the hydrogen bonding distance. In contrast, with Chain A, the distance between the NZ in K26 and OD1 and OD2 in D16 approached approximately 3 Å, indicative of a hydrogen bond. Notably, when the NZ forms a hydrogen bond with OD1, then the hydrogen bond with OD2 is not present (0 - 100 ns & 240 - 300 ns), and when a hydrogen bond is formed with OD2, then the hydrogen bond with OD1 is broken (100 - 240 ns) This suggests that both OD1 and OD2 in Chain A of DAP12 can form hydrogen bonds to NZ in TREM2 but not at the same time.

**Figure 5:**
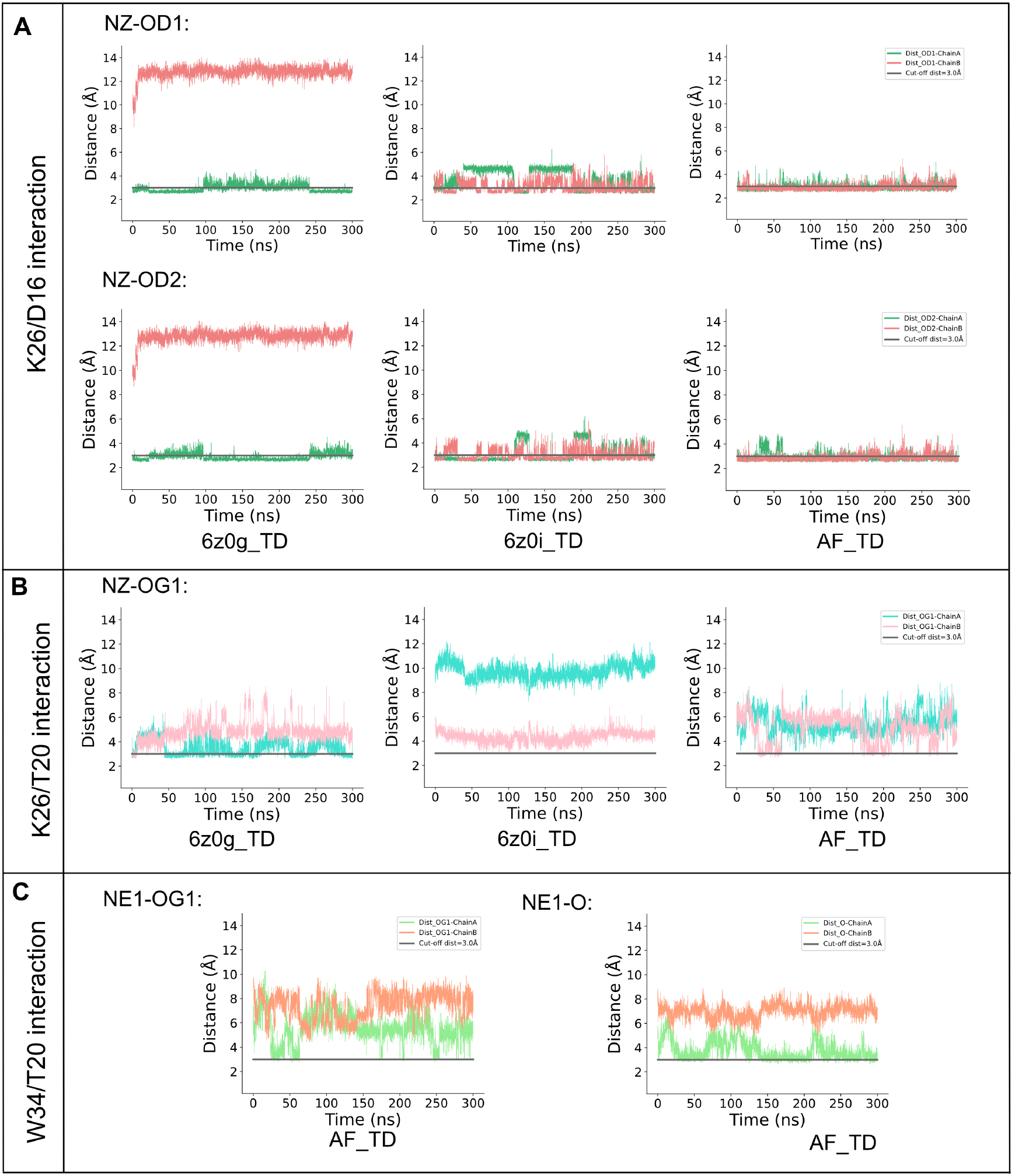
Key hydrogen bond binding residues between TREM2 and DAP12. (A) illustrates the hydrogen bond distances between K26 of TREM2 and D16 of DAP12 in Chain A. The distance for OD1 is represented in green, while that for OD2 is in red. The grey line indicates a cut-off distance of 3 Å. The TD OD1 distance is calculated using NZ (K26) and OD1 (D16), and the TD OD2 distance is calculated using NZ (K26) and OD2 (D16). (B) displays the hydrogen bond distance of K26/T20 in the three systems, calculated using NZ (K26) and OG1 (T20). (C) demonstrates the hydrogen bond distance of W34/T20 in the AF TD system. AF TD OG1 represents the distance between NE1 (W34) and OG1 (T20), while AF TD O represents the distance between NE1 (W34) and O (T20).

In 6z0i TD, again we observe that the hydrogen bonds formed by NZ in TREM2 alternate between the OD1 and OD2 atoms in Chain A of DAP12, where one breaks when the other forms (Figure 5A). For example, the distance between OD1 in Chain A of DAP12 and NZ in TREM2 exceeds 3 Å between 50 and 100 ns of our simulation, while the distance of OD2 in Chain A of DAP12 and NZ in TREM2 was less than 3 Å, indicating a hydrogen bond interaction. This suggests that both OD1 and OD2 can bind to NZ, but only one of OD1 or OD2 can bind to NZ at a specific time. This compensatory mechanism was also observed in Chain B but over a shorter period. Moreover, it was noted that both Chain A and Chain B could form hydrogen bonds in 6z0i TD, while hydrogen bonds were formed with only Chain A in 6z0g TD.

In AF TD, similar hydrogen bonding characteristics were observed as seen in 6z0i TD, where both DAP12 chains could form hydrogen bonds between their OD1 and OD2 atoms with the NZ in the K26 residue of TREM2. Given that 6z0i TD and AF TD systems featured less kinked TREM2 structures, it is inferred that the kinked structure of TREM2 (6z0g TD) results in it only being able to form hydrogen bonds to one chain of DAP12 as opposed to both chains when it is unkinked.

In the second hydrogen bond interaction pair we considered, namely K26/T20, the distance was measured between the NZ atom of the K26 residue in TREM2 and the O atom in the hydroxyl group (OG1, Figure 4E) in the two DAP12 chains. In 6z0i TD, a K26/T20 hydrogen bond occurs between TREM2 and Chain A of DAP12 briefly (e.g., 50 - 100 ns, Figure 5B), but no hydrogen bond is observed with Chain B of DAP12. Meanwhile in the unkinked models (6z0i TD & AF TD), we observe a K26/T20 hydrogen bond only briefly and very transiently in the AF TD model.

In the final hydrogen bond interaction residue pair (Figure 5C), W34/T20, we measured the hydrogen bond distances between the NE1 atom within the W34 residue of TREM2 and the OG1 and O atoms in the T20 residues of DAP12 (Figure 4F). We didn’t find any of these hydrogen bonds in the 6z0g TD or 6z0i TD systems (Figure S10). However in the AF TD model, we found that NE1 in the W34 residue of TREM2 is more commonly found to hydrogen bond to O in the T20 residue of Chain A of DAP12, than with the OG1 atom in the same residue and chain.

### Probabilities of Hydrogen Bonds Formation in Different TREM2/DAP12 Systems

To gain a deeper understanding of the binding mechanism, we calculated the fraction of time that a given hydrogen bond interaction is observed in each of our systems. Table 1 summarises the probabilities for each of the hydrogen bonding pairs that we have identified as being important in the formation of the TREM2/DAP12 complex. In the case of the K26/D16 hydrogen bonds, we found that it is more common to have one formed between the OD2 atom within the D16 residue of DAP12 than with OD1 for all of the different models we studied. As seen previously in Figure 5, the calculated probabilities show that the unkinked models (6z0i TD & AF TD) are able to form K26/D16 hydrogen bonds with both chains of DAP12 in almost equal proportions, whereas the kinked model (6z0g TD) only forms hydrogen bonds with one of the chains of DAP12. Also we see that the K26/D16 hydrogen bonds are much more prevalent in the unkinked models than in the kinked model. The detailed hydrogen bond possibilities of each hydrogen are also shown (Table S1 - S3).

**Table 1:** Hydrogen bond possibilities of TREM2/DAP12 in three systems.

| system | residue | H-bond | Chain A <sup>a</sup> | Chain B <sup>a</sup> |
| --- | --- | --- | --- | --- |
| 6z0g_TD | K26/D16 | NZ-OD1 | 34.74 | 0 |
|  |  | NZ-OD2 | 44.57 | 0 |
|  | K26/T20 | NZ-OG1 | 15.10 | 0.52 |
|  | W34/T20 | NE1-OG1 | 0 | 0 |
|  |  | NE1-O | 0 | 0 |
| 6z0i_TD | K26/D16 | NZ-OD1 | 24.85 | 34.91 |
|  |  | NZ-OD2 | 60.35 | 43.03 |
|  | K26/T20 | NZ-OG1 | 0 | 0 |
|  | W34/T20 | NE1-OG1 | 0 | 0 |
|  |  | NE1-O | 0 | 0 |
| AF_TD | K26/D16 | NZ-OD1 | 32.86 | 33.29 |
|  |  | NZ-OD2 | 38.49 | 34.78 |
|  | K26/T20 | NZ-OG1 | 0 | 1.02 |
|  | W34/T20 | NE1-OG1 | 0.55 | 0 |
|  |  | NE1-O | 9.08 | 0 |

<sup>a</sup>The hydrogen bonds are calculated between TREM2 and DAP 12 Chain A.
<sup>b</sup>The hydrogen bonds are calculated between TREM2 and DAP 12 Chain B.

We also computed the fraction of time that a hydrogen bond is formed between the K26 residue on TREM2 and the T20 residues on DAP12 (Table 1). In the 6z0g TD system, TREM2 exhibits a 15.1% probability of forming a hydrogen bond with Chain A, whereas it is only 0.5% for Chain B. However, in the other two unkinked models, we only see a hydrogen bond formed between these residues a very small fraction of the time (1.0% in the AF TD model).

The fraction of time that the hydrogen bond between the W34 of TREM2 and the T20 residues of DAP12 was observed is also reported in Table 1. We only see this hydrogen bond in AF TD system and again it is only found for a reasonably small amount of the time (9.1% of the time with the O atom in Chain A and 0.6% of the time with the OG1 atom).

In conclusion, our study identified three distinct hydrogen bonding interactions, among which the K26/D16 hydrogen bond is found to be the most significant. This is evident not only because it is present in all three systems but also due to its significantly longer duration compared to K26/T20 or W34/T20 interactions. Notably, in the model in which TREM2 has a kinked structure (6z0g TD), the K26/D16 hydrogen interaction showcases a preference for binding to a specific chain of DAP12, whereas in the unkinked (6z0i TD and AF TD) systems, TREM2 exhibits the capability to bind with both chains of DAP12. Additionally, the W34/T20 hydrogen bond interaction is exclusively observed in AF TD. Meanwhile, the kinked structure (6z0g TD) shows another hydrogen bonding interaction present within the complex, the K26/T20 hydrogen bond interaction.

### Atomic Binding Mechanism of TREM2/DAP12 in Three Systems

We selected a representative snapshot (at *t* = 150 ns) to display the binding interface of TREM2 and DAP12 in the three different systems (Figure 6). In the system with the kinked TREM2 (6z0g TD), the K26 residue of TREM2 binds to the D16 and T20 residues of Chain A of DAP12, forming NZ-HZ2-OD1 and NZ-HZ1-OG1 hydrogen bonds, respectively (Figure 6A). The number of hydrogen bonds (*N_HB_*) is also illustrated in Figure 6A to facilitate a better understanding of the distribution of the various types of hydrogen bonds. Within the 6z0i TD system, hydrogen bonds are exclusively formed between K26 and D16 (Figure 6A). This system forms two hydrogen bonds (NZ-HZ1-OD2 & NZ-HZ3-OD2) during the course of the simulation. In the AF TD system, all three types of hydrogen bonds are detected (Figure 6C). Within the interface, NZ-HZ3-OD2 and NZ-HZ2-OD1 form two of the hydrogen bonds. Additionally, we display the third hydrogen bond NE1-HE1-OG1 in W34/T20 in the interface (illustrated as a grey dashed line) (Figure 6C).

**Figure 6:**
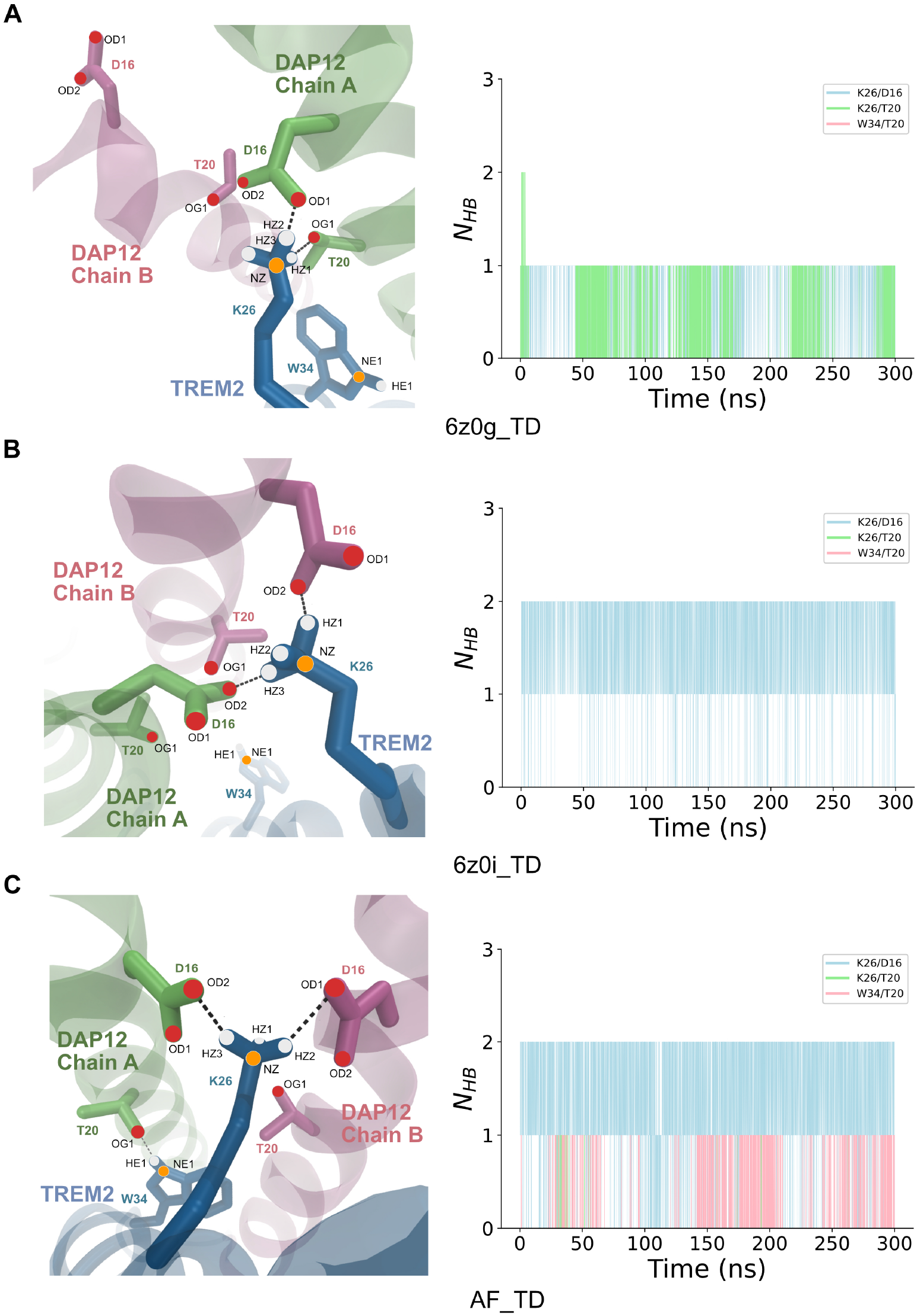
Binding interface of hydrogen bond interactions in three systems. (A) The left picture displays the hydrogen binding interface of 6z0g TD. Black dashed lines represent the observed hydrogen bonds within the timeframe. The right picture showcases the count of hydrogen bonds of K26/D16, K26/T20, and W34/T20, coloured in blue, green, and pink, respectively. (B) illustrates the hydrogen bond interface and the count of hydrogen bonds in 6z0i TD. (C) demonstrates the hydrogen bond interface and the count of hydrogen bonds in AF TD. The grey dashed line indicates the hydrogen bond of W34/T20.

The combined total of hydrogen bonds found in any of the three different models for the TREM2/DAP12 complex is found to be generally two, and in the AF TD model occasionally there is a third hydrogen bond formed (Figures S16A, S16B & S16C).

## Discussion

We understand that TREM2 and DAP12 play crucial roles in Alzheimer’s disease, but the structure of their complex remains elusive to date. Wet lab experiments have highlighted the significance of K26 in TREM2, but specific details have remained mysterious. An interesting comparison in exploring the structure of the TREM2/DAP12 complex was observed in the structure of NKG2C (Natural Killer Cell Group 2)/DAP12,^19^ which shares some commonalities with TREM2/DAP12. However, due to limitations in protein purification, the authors introduced covalent bonds between NKG2C and DAP12, potentially impacting the interaction. Our model precisely utilizes the same sequences in three models and unveils the binding modes of both kinked and unkinked TREM2 structures. Additionally, we accurately determine the hydrogen bonds between TREM2/DAP12 and visualize the changing hydrogen bond patterns. These findings align with previous wet lab discoveries, elucidating the significance of the kink is essential for the complex’s functionality. Remarkably, the predicted structure by AlphaFold2 closely resembles the 6z0i TD structure.

Moreover, we have demonstrated the presence of hydrophobic interactions between TREM2 and DAP12 (Figure 4C). Therefore, we generated contact maps for the hydrophobic interactions between L19/I12, W34/I23, W34/A24, and W34/A27 (Figures S17, S18, S19, and S20, respectively). The AF TD system exhibited the most prevalent hydrophobic interactions among the three systems, 6z0i TD showed the least, and 6z0g TD demonstrated a moderate level of hydrophobic interactions (Figure S21). Considering the interplay of hydrogen bonds, hydrophobic interactions, and salt bridges, we developed an atomic-level TREM2/DAP12 binding model (Figure 7). The snapshot is captured at the 150 ns timeframe in the 6z0i TD system, revealing K26/D16 as the primary contact residue pair capable of forming four hydrogen bonds with NZ in the K26 residue of TREM2 and OD1 and OD2 in the D16 residue in both Chain A and Chain B of DAP12.

**Figure 7:**
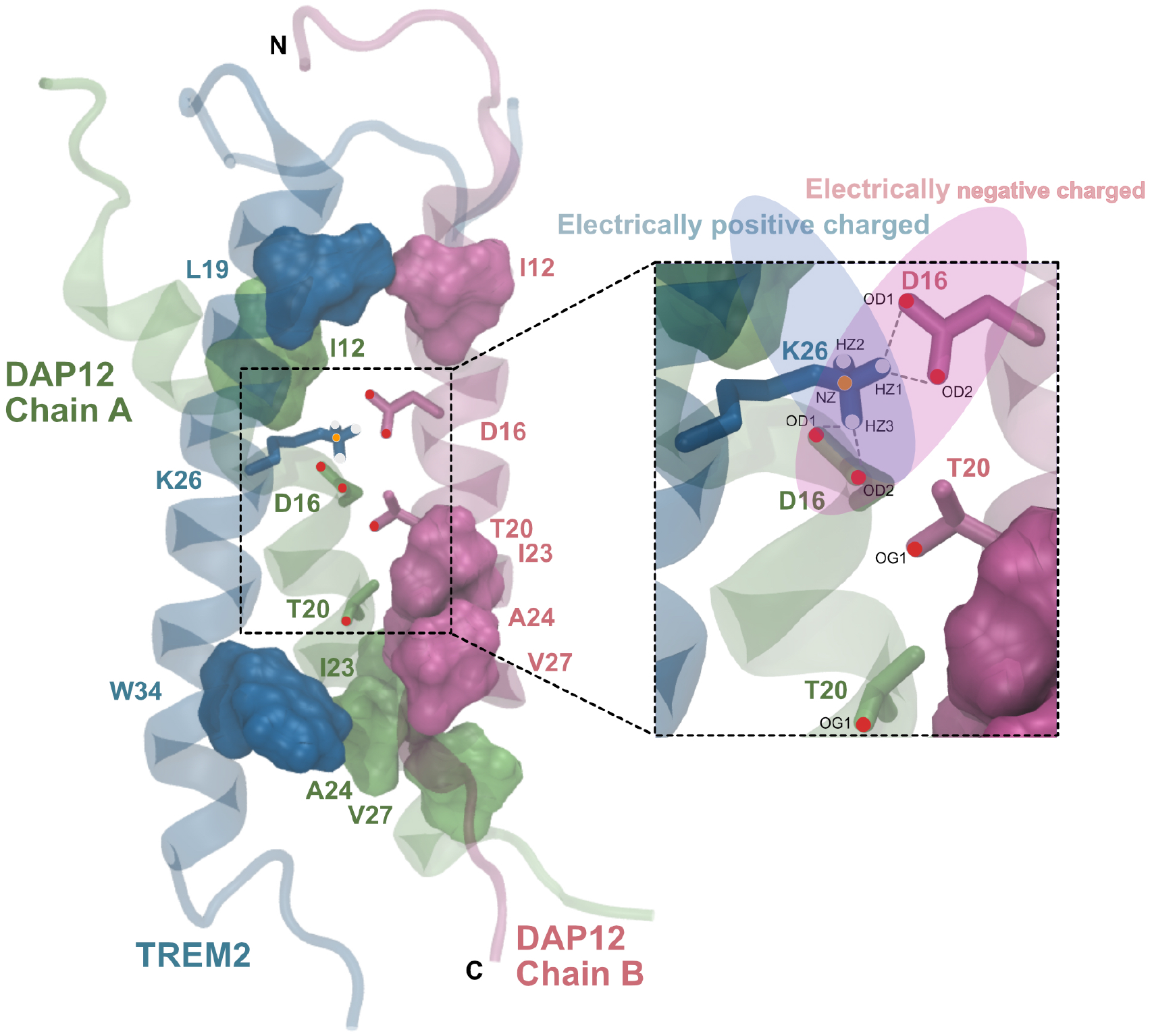
Proposed binding mechanism of TREM2/DAP12. TREM2, DAP12 Chain A, and Chain B are coloured in blue, lime, and mauve, respectively. The blue shadow oval region indicates the positively charged area attributed to K26 in TREM2, while red indicates the negatively charged area attributed to D16 in DAP12, depicting the K26/D16 salt bridge interaction. Grey dashed lines indicate potential hydrogen bonds. Hydrophobbic residue pairs L19/I12, W34/I23, W34/A24, and W34/A27 are shown in surface.

It is notable that in the 6z0i TD system, the hydrogen bond interaction only existed in K26/D16, and the probability of finding the hydrogen bond is relatively large (85.2% and 77.9% for Chain A and Chain B respectively, Table 1). A lower probability of the K26/D16 hydrogen bond existing was found in AF TD (71.4% and 68.1% for Chain A and Chain B, respectively). Also, the AF TD system has a hydrogen bond between the W34 residue in TREM2 and the T20 residues in DAP12 9.1% of the time, and a hydrogen bond between the K26 residue of TREM2 and the T20 residues of DAP12 1.0% of the time. These three types of hydrogen bonds, along with the strong hydrophobic interactions detected in AF TD (Figure S21), contribute to the stability of the AF TD system. In contrast, 6z0i TD exhibits very limited hydrophobic interaction, and only the K26/D16 hydrogen bond interaction has been observed.

We generated representative snapshots illustrating the orientation of hydrophobic residues in three systems (Figure S22). In general, the hydrophobic residues in AF TD predominantly face inward, while those in 6z0i TD exhibit on the outward facing side of the proteins, which allows them to interact with the lipids surrounding them. In our proposed model of the TREM2/DAP12 complex, the receptor TREM2 primarily binds to the DAP12 dimer through hydrogen bonds formed by K26 in TREM2 and D16 in both chains of DAP12 (Figure 7). Additionally, hydrophobic interactions also play significant roles, with K26/T20 and W34/T20 forming two hydrogen bonds that contribute to system stabilization, albeit to a lesser extent than the K26/D16 interaction.

In addition to providing this picture of the stable TREM2/DAP12 complex, our simulations also provide insight into the mechanisms which drive the transition of TREM2 from its unbound (kinked) state to its bound (unkinked) state. The kinked state (6z0g TD) interacts with one chain of DAP12 via a hydrogen bond between its K26 residue and either the D16 residue or the T20 residue on that chain of DAP12. During the binding process, TREM2 undergoes a conformational chain such that it is no longer kinked (6z0i TD & AF TD) and this allows it to interact with both chains of DAP12 and form stable hydrogen bonds between its K26 residue and both of the D16 residues on either chain of DAP12. We also find evidence of hydrophobic interactions that form between the TREM2 and the two chains of DAP12. Therefore, the transitioning from the kinked to unkinked state by TREM2 allows it to form a more stable complex with DAP12.

## Methods

### Systems Built Approach

The 6z0g TD and 6z0i TD complexes were created with HADDOCK^24,25^ and PyMOL using PDB access codes of 6z0g, 6z0i,^26^ and 2l34.^27^ The AF TD were generated with ColabFold^28,29^ v1.5.2 using the same sequence as 6z0g TD and 6z0i TD (the sequence is shown in Table S4). The systems utilized in this article were generated through the use of the CHARMM-GUI Membrane Builder^30^ and the MARTINI Maker^31^ for all-atom (AA) and coarse-grained (CG) simulation respectively. The membrane consists of POPC and cholesterol at a ratio of 80 : 20. The solvent employed in every simulation was water, and to maintain a neutral charge and a 0.15 M salt concentration, Na^+^ and Cl*^−^* ions were added to the system.

### Coarse-grained Molecular Dynamics Simulations

We carried out 200 *µ*s CG simulations to gain stable systems in a POPC/cholesterol lipid bilayer. These CG simulations were using GROMACS^32^ version 2019.2 on the Young cluster. MARTINI22P forcefield^33^ was employed in this system, and each lipid membrane system was first minimized and then equilibrated at the temperature of 310.15 K. The MARTINI22P model groups three to five heavy (non-hydrogen) atoms into each interacting bead, with different bead types representing various combinations of grouped atoms and chemical environments. There are four groups of bead type: charged, polar, non-polar, and polar, each with five distinct types distinguished by their polar affinity or tendency to form hydrogen bonds in the MARTINI22P forcefield. The details of the CG simulation of different TREM2/DAP12 systems are shown in Table 2. We carried out three replicas of each system.

**Table 2:** Details of each of the CG simulation systems.

| system | $n_{\text{atoms}}$ | $n_{\text{lipids}}$ | $n_{\text{waters}}$ | $n_{\text{ions}}$ | final box size (nm) |
| --- | --- | --- | --- | --- | --- |
| 6z0g_TD<sup>a</sup> | 9287 | 210 | 5463 | 125 | $100.6 \times 100.6 \times 112.0$ |
| 6z0i_TD | 9336 | 210 | 5510 | 127 | $100.6 \times 100.6 \times 113.0$ |
| AF_TD | 11320 | 315 | 7398 | 167 | $101.0 \times 101.0 \times 134.7$ |

### All-atom Molecular Dynamics Simulations

After 200 *µ*s of CG simulations, we converted the CG simulation to all-atom simulations using CHARMM-GUI Martini to All-atom Converter,^30,34^ and ran another 300 ns. We utilized GROMACS 2020.3 for these AA simulations on the CREATE cluster. Our AA simulations utilized the same POPC/cholesterol bilayer, with lipids modelled using the CHARMM36 forcefield^35,36^ and CHARMM modified TIP3P water model.^37^ To ensure accurate results, each lipid membrane system was first minimized and then equilibrated at a temperature of 310.15 K and a pressure of 1 bar according to the simulation protocol prescribed by CHARMM-GUI.^37,38^ The same simulation procedure was applied to all systems. It commenced with an initial energy minimization step using the steepest descent algorithm, followed by a 125 picosecond (ps) simulation within the NVT ensemble, where the Nos‘–Hoover thermostat was employed to regulate the temperature (set at 310.15 K) with a time step of 1 femtosecond (fs). Subsequently, 1 *µ*s production simulations were executed within the NPT ensemble, utilizing both the Nos‘–Hoover thermostat and the Parrinello–Rahman barostat to maintain a constant temperature of 310.15 K and a pressure of 1 atmosphere (atm). Throughout these simulations, hydrogen-containing bonds were kept fixed using the LINCS algorithm.^39^ Periodic boundary conditions were employed across all dimensions throughout all simulations, with a time step of 1 fs utilized for both equilibration and production simulations. The cutoff used for the calculation of Coulombic and Lennard-Jones (LJ) forces was 12 Å, and a switching function reduced the LJ interactions to zero from an inner cutoff of 10 Å. Table 3 displays the AA simulation details of the TREM2/DAP12 complex. Three replicas were performed.

**Table 3:** Details of each of the AA simulation systems.

| system | $n_{\text{atoms}}$ | $n_{\text{lipids}}$ | $n_{\text{waters}}$ | $n_{\text{ions}}$ | final box size (nm) |
| --- | --- | --- | --- | --- | --- |
| 6z0g_TD | 77286 | 220 | 16214 | 87 | $85.7 \times 85.7 \times 112.3$ |
| 6z0i_TD | 78446 | 220 | 16600 | 89 | $85.6 \times 85.6 \times 115.2$ |
| AF_TD | 94204 | 225 | 21640 | 117 | $86.0 \times 86.0 \times 136.0$ |

### Molecular Dynamics Simulation Analysis

The analysis of all data was performed using in-house Python (3.10) scripts, which make wide use of the MDAnalysis package ^40,41^ and visualised using Matplotlib.^42^ Furthermore, all simulations conducted in this study were performed under periodic boundary conditions, with the resulting trajectories viewed through the utilization of VMD^43^ 1.9.3. The kinked angles were calculated using the C*_α_* atoms in TREM2, specifically utilizing the residues A20 and L24 on TREM2 as the first vector and residues A32 and A36 as the second vector. The contact map presented in Figure 3 was generated using the C*_α_* atoms between TREM2 and DAP12 with a cut-off distance set at 11 Å. The hydrogen bond possibility was computed utilizing the Hydrogen Bond Analysis^44^ function within MDAnalysis and was executed using an in-house Python script. The cut-off distance for the hydrogen bond is set to 3 Å and the cut-off angle is set to 150 °.

## Acknowledgement

We thank Dr. R.L.D. Castro at King’s College London for her computational assistance. We are grateful to the UK Materials and Molecular Modelling Hub for computational resources, which is partially funded by EPSRC (EP/T022213/1, EP/W032260/1 and EP/P020194/1). We also appreciate the King’s Computational Research, Engineering and Technology Environment (CREATE) at King’s College London for providing us access to computational resources. Z.Z. acknowledges receiving PhD studentships from King’s College London and China Scholarship Council.

## Supporting Information Available

All study data are included in the article and/or SI Appendix.

## Contribution

Z.Z., M.U., and C.D.L. designed research; Z.Z performed research; Z.Z., M.U., and C.D.L. analyzed data; Z.Z and C.D.L wrote the paper.

## References

(1) Association, A.; others 2016 Alzheimer’s disease facts and figures. Alzheimer’s & Dementia 2016, 12, 459–509.

(2) Drouin, E.; Drouin, G. The first report of Alzheimer’s disease. The Lancet Neurology 2017, 16, 687.

(3) Carmona, S.; Zahs, K.; Wu, E.; Dakin, K.; Bras, J.; Guerreiro, R. The role of TREM2 in Alzheimer’s disease and other neurodegenerative disorders. The Lancet Neurology 2018, 17, 721–730.

(4) Kidd, M. Alzheimer’s disease—an electron microscopical study. Brain 1964, 87, 307–320.

(5) Golde, T. E.; Estus, S.; Younkin, L. H.; Selkoe, D. J.; Younkin, S. G. Processing of the amyloid protein precursor to potentially amyloidogenic derivatives. Science 1992, 255, 728–730.

(6) Guerreiro, R.; Wojtas, A.; Bras, J.; Carrasquillo, M.; Rogaeva, E.; Majounie, E.; Cruchaga, C.; Sassi, C.; Kauwe, J. S.; Younkin, S.; others TREM2 variants in Alzheimer’s disease. New England Journal of Medicine 2013, 368, 117–127.

(7) Jonsson, T.; Stefansson, H.; Steinberg, S.; Jonsdottir, I.; Jonsson, P. V.; Snaedal, J.; Bjornsson, S.; Huttenlocher, J.; Levey, A. I.; Lah, J. J.; others Variant of TREM2 associated with the risk of Alzheimer’s disease. New England Journal of Medicine 2013, 368, 107–116.

(8) Müller, U. C.; Deller, T.; Korte, M. Not just amyloid: physiological functions of the amyloid precursor protein family. Nature Reviews Neuroscience 2017, 18, 281–298.

(9) Hamerman, J. A.; Jarjoura, J. R.; Humphrey, M. B.; Nakamura, M. C.; Seaman, W. E.; Lanier, L. L. Cutting edge: inhibition of TLR and FcR responses in macrophages by triggering receptor expressed on myeloid cells (TREM)-2 and DAP12. The Journal of Immunology 2006, 177, 2051–2055.

(10) Peng, Q.; Malhotra, S.; Torchia, J. A.; Kerr, W. G.; Coggeshall, K. M.; Humphrey, M. B. TREM2-and DAP12-dependent activation of PI3K requires DAP10 and is inhibited by SHIP1. Science signaling 2010, 3, ra38–ra38.

(11) Kober, D. L.; Brett, T. J. TREM2-ligand interactions in health and disease. Journal of molecular biology 2017, 429, 1607–1629.

(12) Ulland, T. K.; Song, W. M.; Huang, S. C.-C.; Ulrich, J. D.; Sergushichev, A.; Beatty, W. L.; Loboda, A. A.; Zhou, Y.; Cairns, N. J.; Kambal, A.; others TREM2 maintains microglial metabolic fitness in Alzheimer’s disease. Cell 2017, 170, 649–663.

(13) Zhou, R.; Yang, G.; Guo, X.; Zhou, Q.; Lei, J.; Shi, Y. Recognition of the amyloid precursor protein by human *γ*-secretase. Science 2019, 363, eaaw0930.

(14) Thornton, P.; Sevalle, J.; Deery, M. J.; Fraser, G.; Zhou, Y.; Ståhl, S.; Franssen, E. H.; Dodd, R. B.; Qamar, S.; Gomez Perez-Nievas, B.; others TREM 2 shedding by cleavage at the H157-S158 bond is accelerated for the Alzheimer’s disease-associated H157Y variant. EMBO molecular medicine 2017, 9, 1366–1378.

(15) Feuerbach, D.; Schindler, P.; Barske, C.; Joller, S.; Beng-Louka, E.; Worringer, K. A.; Kommineni, S.; Kaykas, A.; Ho, D. J.; Ye, C.; others ADAM17 is the main sheddase for the generation of human triggering receptor expressed in myeloid cells (hTREM2) ectodomain and cleaves TREM2 after Histidine 157. Neuroscience letters 2017, 660, 109–114.

(16) Schlepckow, K.; Kleinberger, G.; Fukumori, A.; Feederle, R.; Lichtenthaler, S. F.; Steiner, H.; Haass, C. An Alzheimer-associated TREM2 variant occurs at the ADAM cleavage site and affects shedding and phagocytic function. EMBO molecular medicine 2017, 9, 1356–1365.

(17) Yang, J.; Fu, Z.; Zhang, X.; Xiong, M.; Meng, L.; Zhang, Z. TREM2 ectodomain and its soluble form in Alzheimer’s disease. Journal of neuroinflammation 2020, 17, 1–12.

(18) Colonna, M.; Wang, Y. TREM2 variants: new keys to decipher Alzheimer disease pathogenesis. Nature Reviews Neuroscience 2016, 17, 201–207.

(19) Call, M. E.; Wucherpfennig, K. W.; Chou, J. J. The structural basis for intramembrane assembly of an activating immunoreceptor complex. Nature immunology 2010, 11, 1023–1029.

(20) Steiner, A.; Schlepckow, K.; Brunner, B.; Steiner, H.; Haass, C.; Hagn, F. *γ*-Secretase cleavage of the Alzheimer risk factor TREM 2 is determined by its intrinsic structural dynamics. The EMBO journal 2020, 39, e104247.

(21) Sinha, S.; Tam, B.; Wang, S. M. Applications of Molecular Dynamics Simulation in Protein Study. Membranes 2022, 12, 844.

(22) Berman, H. M.; Westbrook, J.; Feng, Z.; Gilliland, G.; Bhat, T. N.; Weissig, H.; Shindyalov, I. N.; Bourne, P. E. The protein data bank. Nucleic acids research 2000, 28, 235–242.

(23) Jumper, J.; Evans, R.; Pritzel, A.; Green, T.; Figurnov, M.; Ronneberger, O.; Tunyasuvunakool, K.; Bates, R.; ΁ídek, A.; Potapenko, A.; others Highly accurate protein structure prediction with AlphaFold. Nature 2021, 596, 583–589.

(24) Dominguez, C.; Boelens, R.; Bonvin, A. M. HADDOCK: a protein-protein docking approach based on biochemical or biophysical information. Journal of the American Chemical Society 2003, 125, 1731–1737.

(25) Van Zundert, G.; Rodrigues, J.; Trellet, M.; Schmitz, C.; Kastritis, P.; Karaca, E.; Melquiond, A.; van Dijk, M.; De Vries, S.; Bonvin, A. The HADDOCK2. 2 web server: user-friendly integrative modeling of biomolecular complexes. Journal of molecular biology 2016, 428, 720–725.

(26) Steiner, A.; Schlepckow, K.; Brunner, B.; Steiner, H.; Haass, C.; Hagn, F. *γ*-Secretase cleavage of the Alzheimer risk factor 2 is determined by its intrinsic structural dynamics. The EMBO Journal 2020, 39 .

(27) Call, M. E.; Wucherpfennig, K. W.; Chou, J. J. The structural basis for intramembrane assembly of an activating immunoreceptor complex. Nature Immunology 2010, 11, 1023–1029.

(28) Mirdita, M.; Schütze, K.; Moriwaki, Y.; Heo, L.; Ovchinnikov, S.; Steinegger, M. ColabFold: making protein folding accessible to all. Nature Methods 2022, 19, 679–682.

(29) Jumper, J. et al. Highly accurate protein structure prediction with AlphaFold. Nature 2021, 596, 583–589.

(30) Jo, S.; Kim, T.; Iyer, V. G.; Im, W. CHARMM-GUI: a web-based graphical user interface for CHARMM. Journal of computational chemistry 2008, 29, 1859–1865.

(31) Siewert, J. M.; others The MARTINI force field: coarse grained model for biomolecular simulations. The Journal of Physical Chemistry B 2007, 111, 7812–7824.

(32) Van Der Spoel, D.; Lindahl, E.; Hess, B.; Groenhof, G.; Mark, A. E.; Berendsen, H. J. GROMACS: fast, flexible, and free. Journal of computational chemistry 2005, 26, 1701–1718.

(33) Qi, Y.; Inǵolfsson, H. I.; Cheng, X.; Lee, J.; Marrink, S. J.; Im, W. CHARMM-GUI martini maker for coarse-grained simulations with the martini force field. Journal of chemical theory and computation 2015, 11, 4486–4494.

(34) Qi, Y.; Cheng, X.; Han, W.; Jo, S.; Schulten, K.; Im, W. CHARMM-GUI PACE CG Builder for solution, micelle, and bilayer coarse-grained simulations. Journal of chemical information and modeling 2014, 54, 1003–1009.

(35) Guvench, O.; Mallajosyula, S. S.; Raman, E. P.; Hatcher, E.; Vanommeslaeghe, K.; Foster, T. J.; Jamison, F. W.; MacKerell Jr, A. D. CHARMM additive all-atom force field for carbohydrate derivatives and its utility in polysaccharide and carbohydrate– protein modeling. Journal of chemical theory and computation 2011, 7, 3162–3180.

(36) Klauda, J. B.; Venable, R. M.; Freites, J. A.; O’Connor, J. W.; Tobias, D. J.; Mondragon-Ramirez, C.; Vorobyov, I.; MacKerell Jr, A. D.; Pastor, R. W. Update of the CHARMM all-atom additive force field for lipids: validation on six lipid types. The journal of physical chemistry B 2010, 114, 7830–7843.

(37) MacKerell Jr, A. D.; Bashford, D.; Bellott, M.; Dunbrack Jr, R. L.; Evanseck, J. D.; Field, M. J.; Fischer, S.; Gao, J.; Guo, H.; Ha, S.; others All-atom empirical potential for molecular modeling and dynamics studies of proteins. The journal of physical chemistry B 1998, 102, 3586–3616.

(38) Best, R. B.; Zhu, X.; Shim, J.; Lopes, P. E.; Mittal, J.; Feig, M.; MacKerell Jr, A. D. Optimization of the additive CHARMM all-atom protein force field targeting improved sampling of the backbone *ϕ*, *ψ* and side-chain *χ*1 and *χ*2 dihedral angles. Journal of chemical theory and computation 2012, 8, 3257–3273.

(39) Hess, B.; Bekker, H.; Berendsen, H. J.; Fraaije, J. G. LINCS: A linear constraint solver for molecular simulations. Journal of computational chemistry 1997, 18, 1463–1472.

(40) Gowers, R. J.; Linke, M.; Barnoud, J.; Reddy, T. J.; Melo, M. N.; Seyler, S. L.; Domanski, J.; Dotson, D. L.; Buchoux, S.; Kenney, I. M.; others MDAnalysis: a Python package for the rapid analysis of molecular dynamics simulations. Proceedings of the 15th python in science conference. 2016; p 105.

(41) Michaud-Agrawal, N.; Denning, E. J.; Woolf, T. B.; Beckstein, O. MDAnalysis: a toolkit for the analysis of molecular dynamics simulations. Journal of computational chemistry 2011, 32, 2319–2327.

(42) Hunter, J. D. Matplotlib: A 2D graphics environment. Computing in Science & Engineering 2007, 9, 90–95.

(43) Humphrey, W.; Dalke, A.; Schulten, K. VMD: visual molecular dynamics. Journal of molecular graphics 1996, 14, 33–38.

(44) Smith, P.; Ziolek, R. M.; Gazzarrini, E.; Owen, D. M.; Lorenz, C. D. On the interaction of hyaluronic acid with synovial fluid lipid membranes. Physical Chemistry Chemical Physics 2019, 21, 9845–9857.

