## Supplemental Data 1 for "Understanding the binding mechanisms of Alzheimer risk factor TREM2 in lipid bilayer using multi-scale molecular dynamics simulations"

**This PDF file includes:**

Supporting text
Figs. S1 to S22
Tables S1 to S4

### Supporting Information Text

**Stability check of the three systems.** Fig. S1 depicts the total potential contacts between TREM2 and DAP12. The data spans four different timescales (25 - 75 ns, 100 - 150 ns, 175 - 225 ns, and 250 - 300 ns) within a 300 ns all-atom (AA) simulation, comprising 50 ns simulation intervals. Potential contacts are assessed based on $C_\alpha$ atom distances between TREM2 and DAP12 that are less than 11.0 Å. In the case of 6z0g_TD, the potential contacts for Chain A occur between 100 - 150 ns, while those for Chain B span 70 - 150 ns. For 6z0i_TD, the potential contacts for Chain A remain relatively stable throughout the simulation, reaching around 100 at the conclusion. In contrast, the potential contacts for Chain B fluctuate significantly between 25 - 75 ns and stabilize toward the end of the simulation, also reaching a count of about 100. In the case of AF_TD, the potential contacts exhibit a slight decrease and eventually reach about 100 in both chains.

We used the potential contact maps to assess the stability of the three systems. In the 6z0g_TD system (Fig. S2), the potential contact maps for both Chain A (Fig. S2A) and Chain B (Fig. S2B) of DAP12 exhibit a high degree of similarity, suggesting a stable state in the complex.

In the 6z0i_TD system, the potential contact maps for Chain A (Fig. S3A) of DAP12 are largely similar. However, minor differences are noticed in the 175 - 225 ns period for 6z0o_TD Chain B (Fig. S3B). These discrepancies are observed in the N-terminal regions of the TREM2/DAP12 trimer, known for containing unstructured loops. Our focus on the $\alpha$-helix region indicates acceptable consistency in the middle region contact maps across all three contact maps.

As for the AF_TD system, the potential contact maps exhibit consistency across the 25 - 75 ns, 100 - 150 ns, and 175 - 225 ns periods (Fig. S4). However, in the 250 - 300 ns period, a decrease in contacts is noted in both chains within the N-terminal region of TREM2/DAP12, while the middle region interactions remain stable. These observations indicate a stable interaction within the TREM2 and DAP12 transmembrane domain (TMD) post the coarse-grained simulations.

**Confidence of the TREM2/DAP12 complex generated by AlphaFold2.** Fig S5 shows the confidence of the AlplaFold2 predicted structures.

**Contact maps of TREM2/DAP12 complex.** Fig. S6 illustrates the contact maps of TREM2/DAP12 using a cutoff distance of 3 Å for all atoms, including side chain atoms. Meanwhile, Fig. S7 depicts the regenerated potential contact maps of the TREM2/DAP12 complex. It uses an indexing system where a dot is placed in the contact map if the contact time exceeds 50 % of the total time.

**Contact maps of key residues in TD complex.** Fig. S8 - S15 display the contact maps of key residue pairs, comprising K26/D16, K26/T20, W34/T20, L19/I12, W34/I23, W34/A24, W34/V27, and W38/Y28, respectively.

**Detailed hydrogen bond possibilities of the TREM2/DAP12.** Table S1 - S3 shows the hydrogen bond possibilities.

**Mergerd hydrogen bond interaction.** Fig. S16 shows the merged hydrogen bond interaction.

**Hydrophobic interaction of key residues.** Fig. S17, S18, S19, and S20 display the residue pairs involved in hydrophobic interactions, namely L19/I12, W23/I23, W34/A24, and W34/V27, respectively. Fig. S21 illustrates the distance between these residue pairs to determine hydrophobic interactions. In this analysis, a hydrophobic interaction is defined when the distance between any carbon atoms within a particular residue and another residue is below 5 Å. Fig. S22 shows the facing direction of hydrophobic residues in three systems. The snapshots were captured in 150 ns.

**Sequence of three systems.** Table S4 shows the amino acid sequence of our built systems.

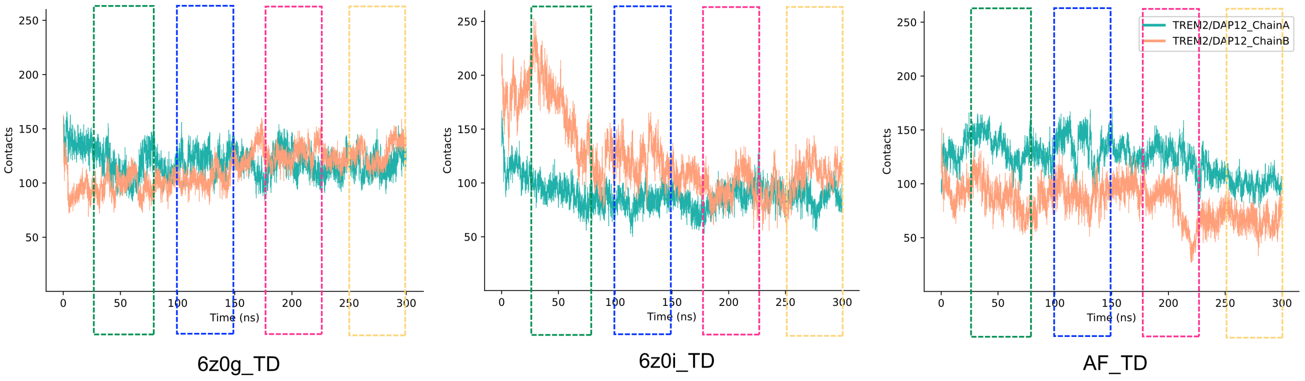


**Fig. S1.** Total contacts across the three systems. Distinctly coloured rectangles indicate various time scales: green, blue, red, and yellow respectively represent 25 - 75 ns, 100 - 150 ns, 175 - 225 ns, and 250 - 300 ns. Contacts are colour-labelled according to the different chains within DAP12; sea-green designates the interaction of TREM2 and DAP12 Chain A, while orange represents the interaction of TREM2 and DAP12 Chain B.

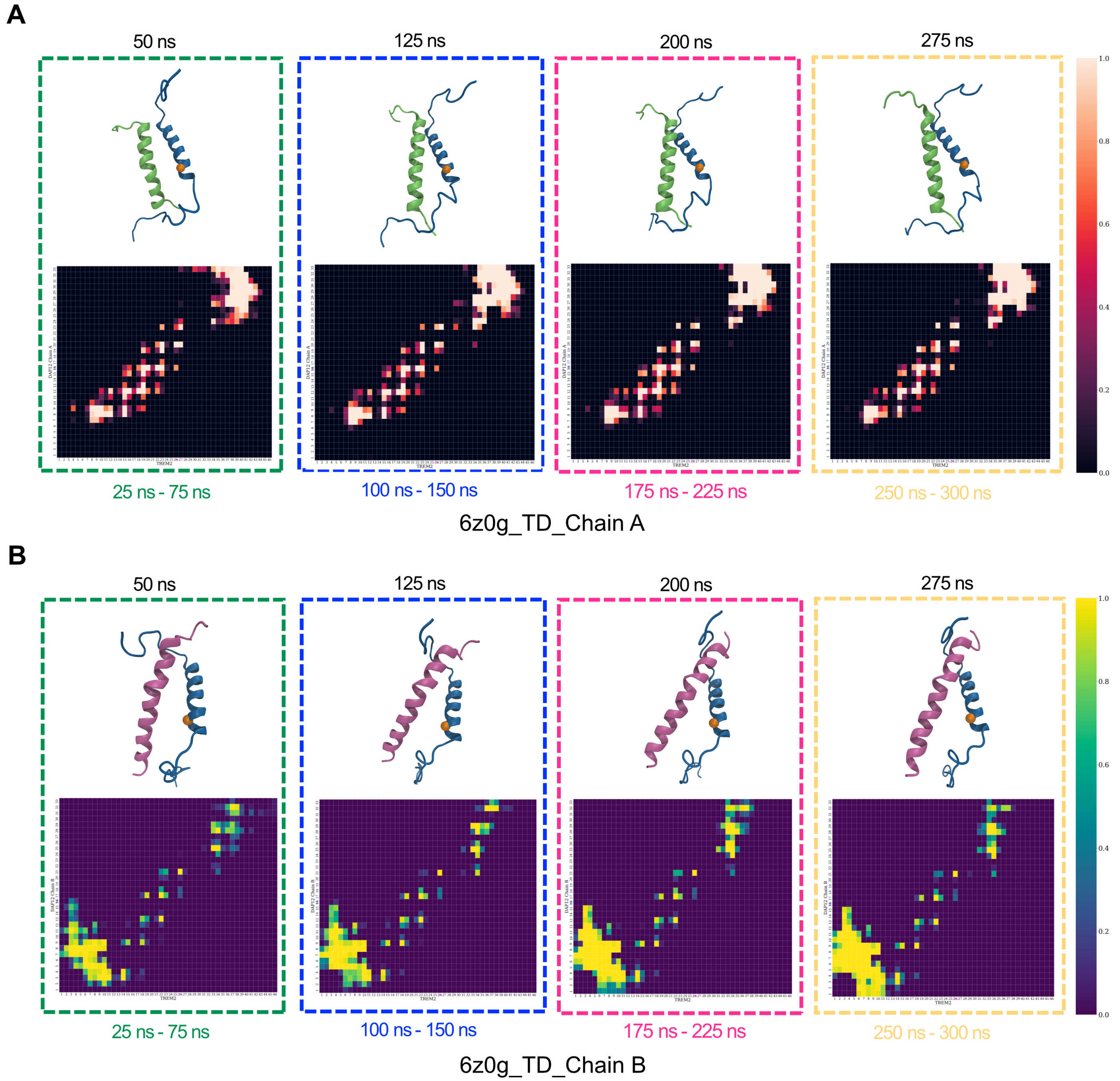


**Fig. S2.** 3D structure snapshots and potential contact maps of 6z0g_TD. (A) The four 3D structures were captured at different times (50 ns, 125 ns, 200 ns, 275 ns) and are represented by differently coloured rectangles, consistent with Fig. S1. The blue-coloured $\alpha$-helix represents TREM2, while the green-coloured $\alpha$-helix shows DAP12 Chain A. The orange sphere denotes the C-$\alpha$ atom of K26 in TREM2. The potential contact maps depict normalized contact numbers, with black indicating 0 and white indicating 1. These contact maps were calculated over four different timescales (25 - 75 ns, 100 - 150 ns, 175 - 225 ns, and 250 - 300 ns), consistent with Fig. S1. (B) In the 3D structure screenshots, the blue-coloured $\alpha$-helix represents TREM2, while the red-coloured $\alpha$-helix shows DAP12 Chain B. The contact maps were normalized to 0 and 1, represented by deep blue and yellow, respectively.

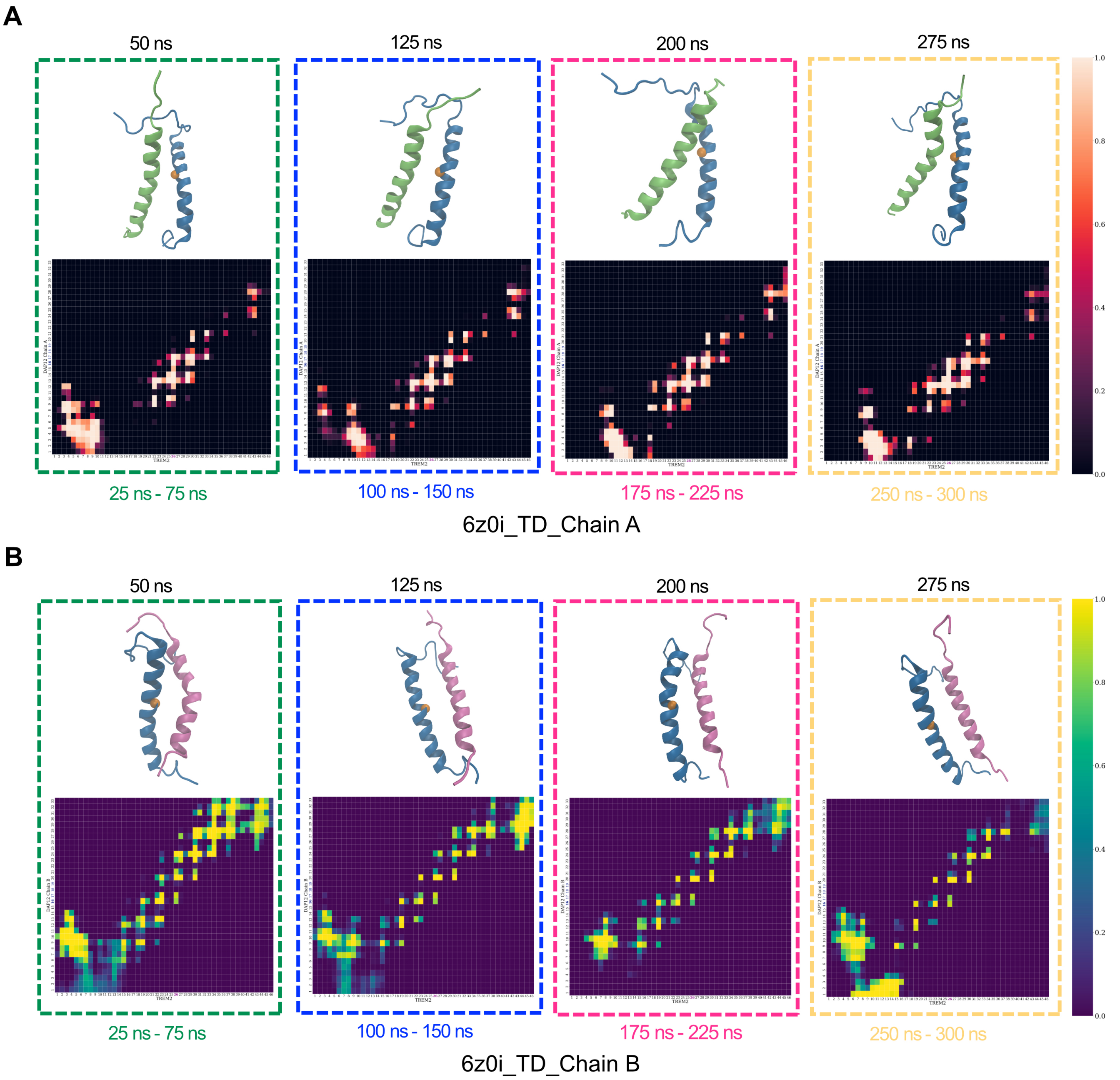


**Fig. S3.** 3D structure screenshots and potential contact maps of 6z0i_TD. (A) and (B) are consistent with Figure S2.

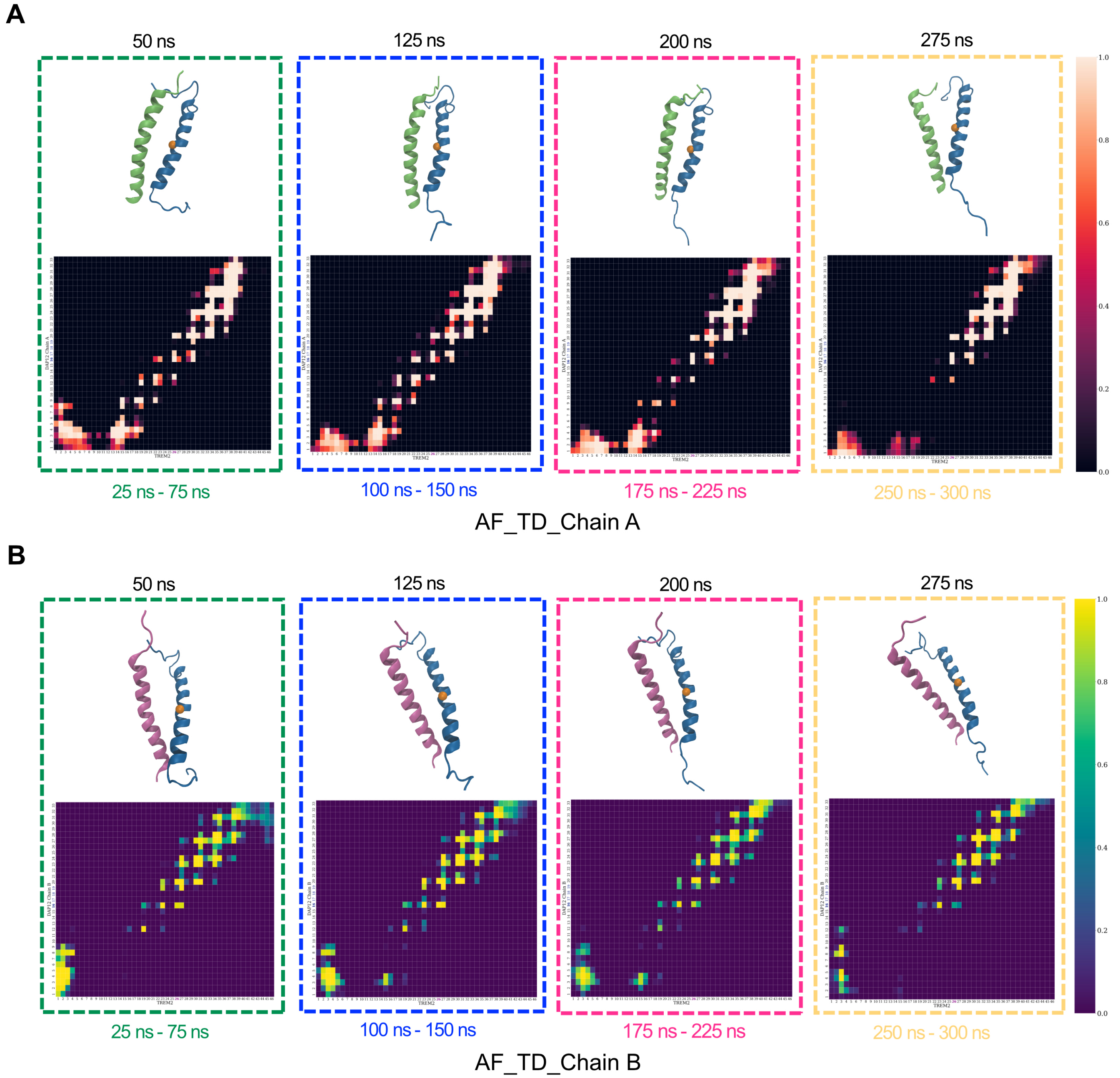


**Fig. S4.** 3D structure screenshots and potential contact maps of AF_TD. (A) and (B) are consistent with Fig. S2.

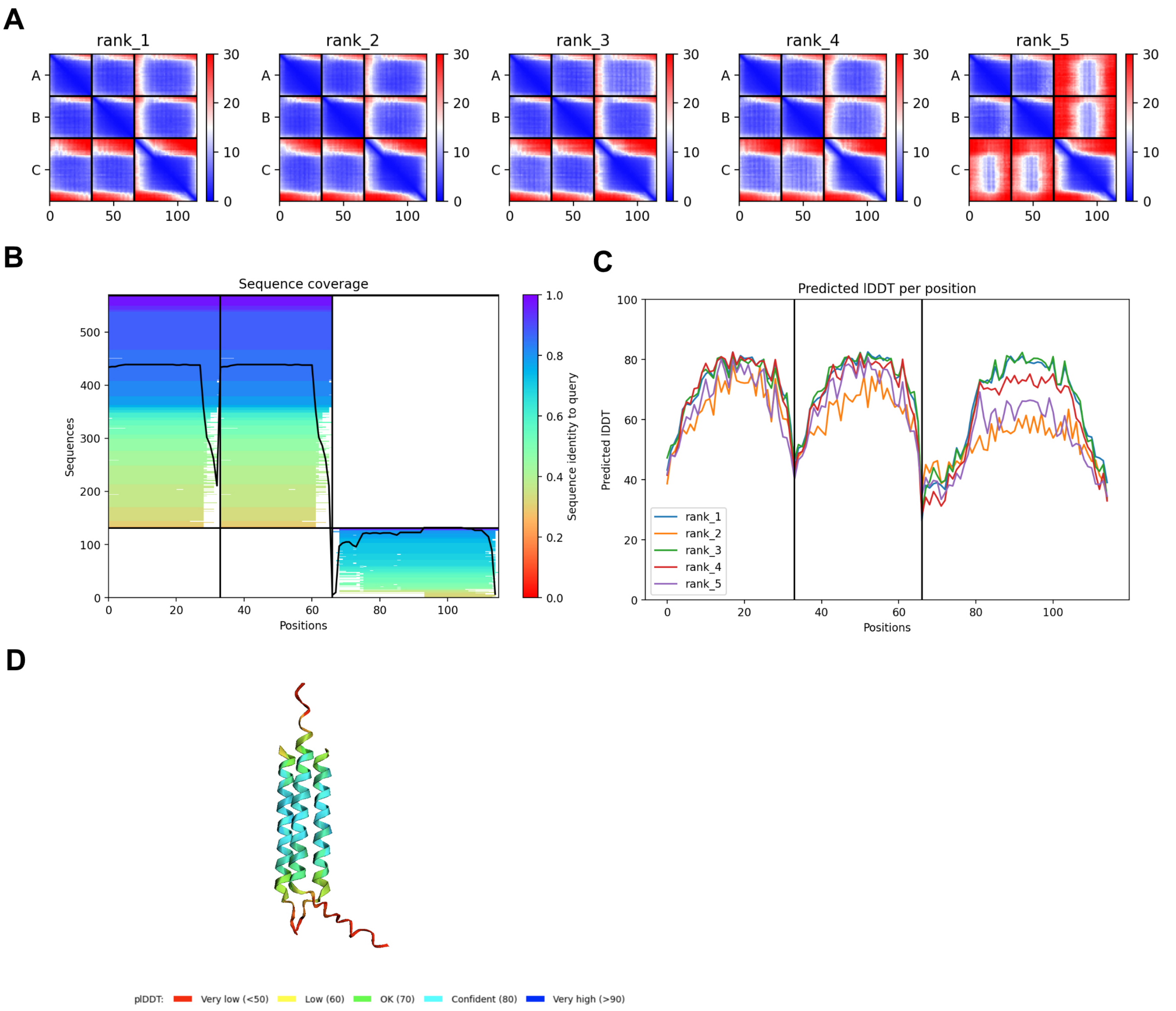


**Fig. S5.** Confidence of AlphaFold2 predicted structures. (A) PAE plots featuring two domains with low conformation confidence. (B) The multiple sequence alignment is summarised as a heatmap. The heat map representation of the MSA indicates all sequences mapped to the input sequences. The colour scale indicates the identity score, and sequences are ordered from top (largest identity) to bottom (lowest identity). White regions are not covered, which occurs with sub-sequence entries in the database. The black line qualifies the relative coverage of the sequence with respect to the total number of aligned sequences. (C) The predicted LDDT per residue for the 5 models was obtained after an AlphaFold2 job. (D) The predicted 3D structure of AlphaFold2 is coloured in predicted LDDT.

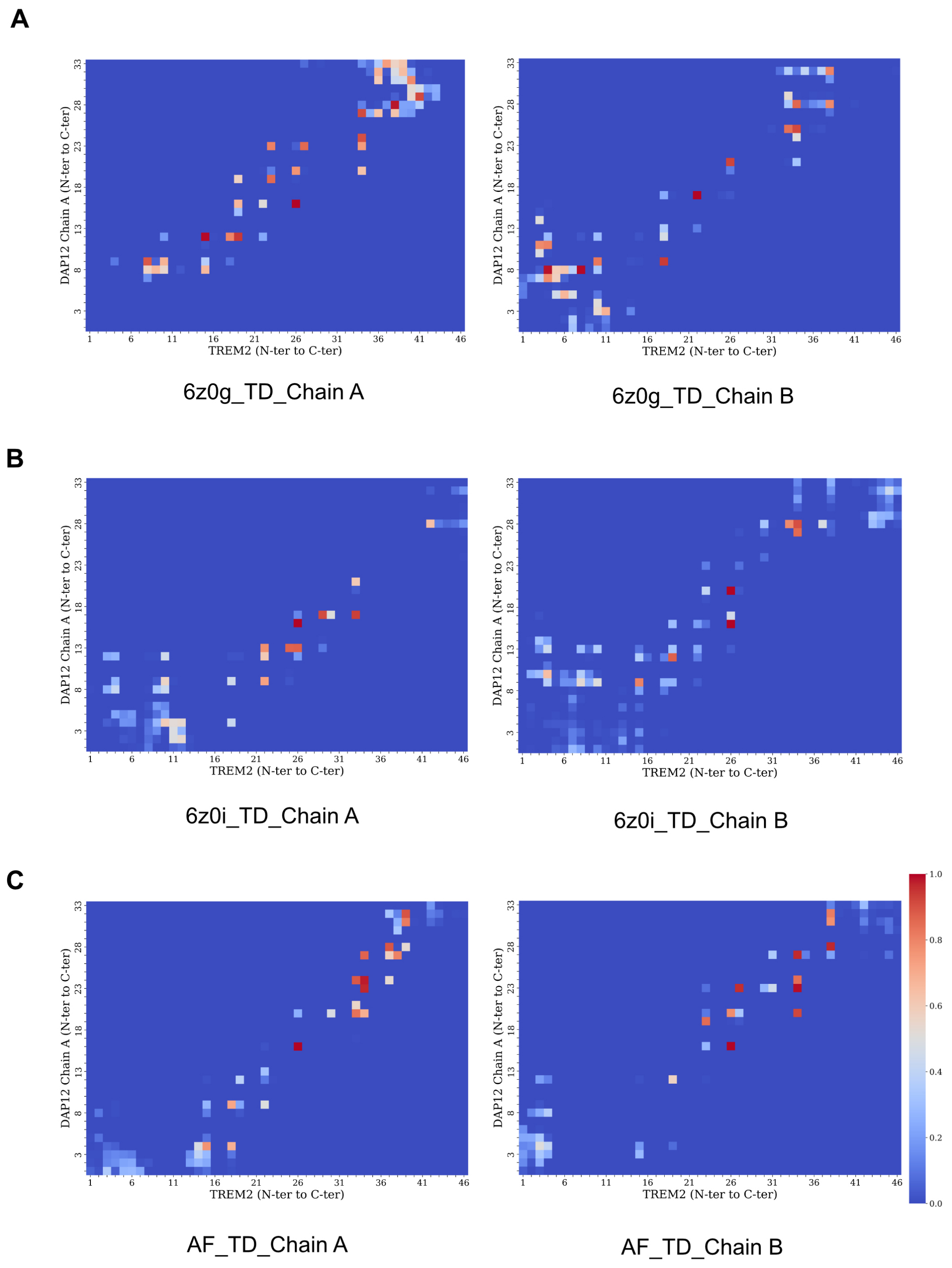


**Fig. S6.** Contact maps of TREM2/DAP12 complex.

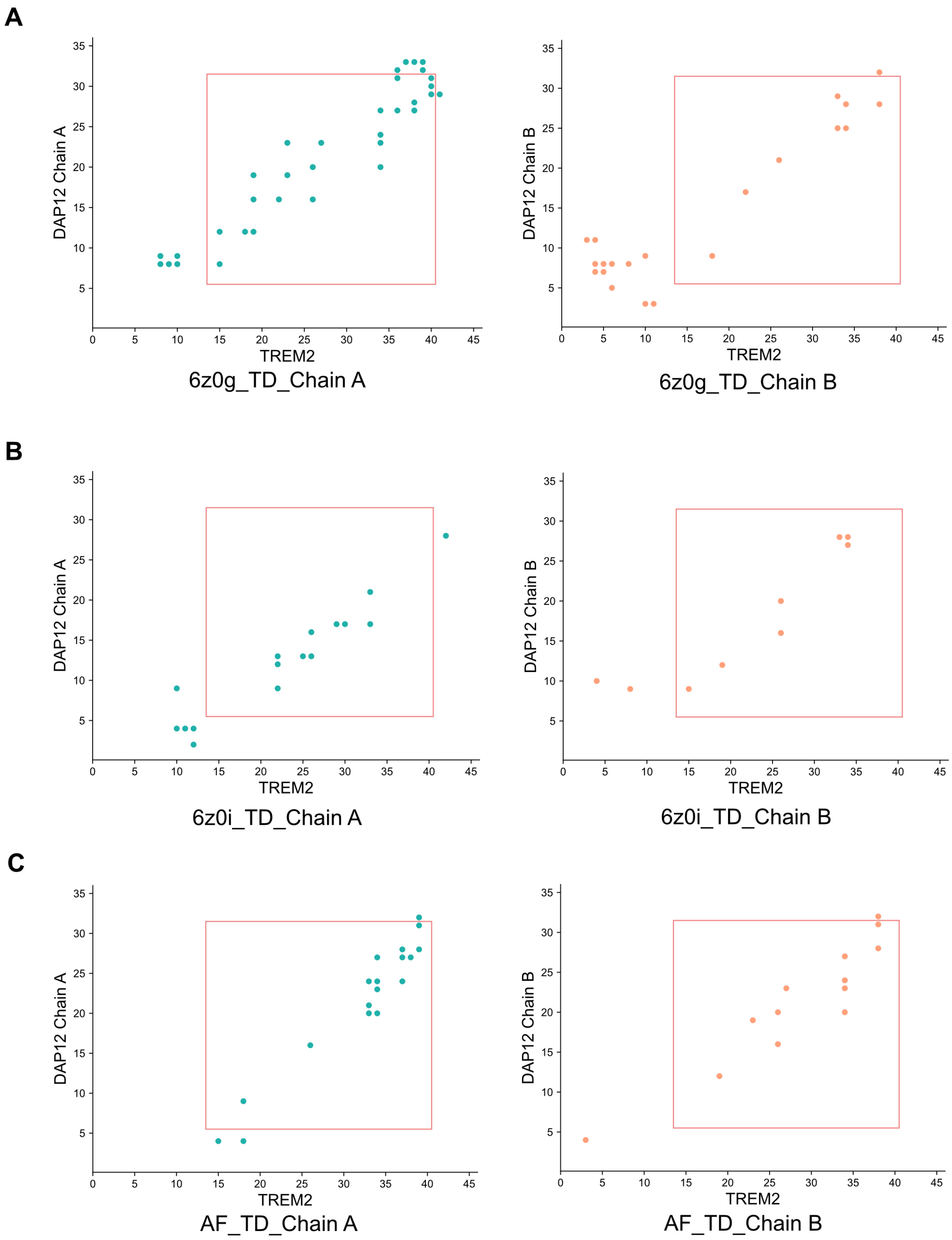


**Fig. S7.** Revised potential contact maps. These calculated potential contact maps were generated using 300 ns trajectory data, and the distance between C$\alpha$ atoms was below 11 Å for no less than 90% of the whole simulation time. (A) shows the calculated contact maps of 6z0g_TD. The contacts between DAP12_Chain A and TREM2 are in green, and between DAP12_Chain B and TREM2 are in orange. The red rectangle represents the $\alpha$-helix, and residues in the rectangle indicate the residues in the $\alpha$-helix, while those outside are in the loop region. (B) shows the calculated contact maps of 6z0i_TD. (C) shows the calculated contact maps of AF_TD.

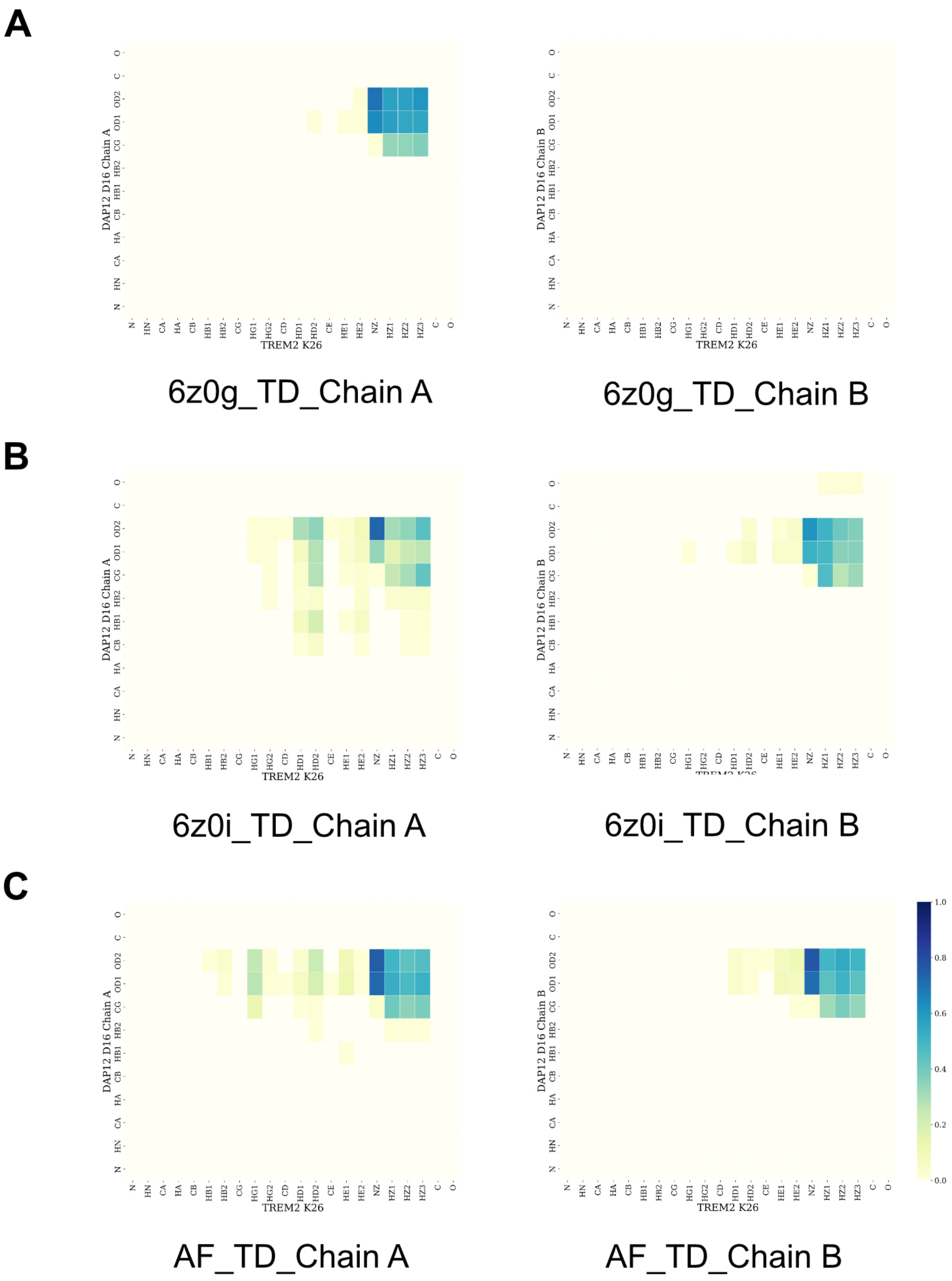


**Fig. S8.** Contact maps of L19/I12 residues.

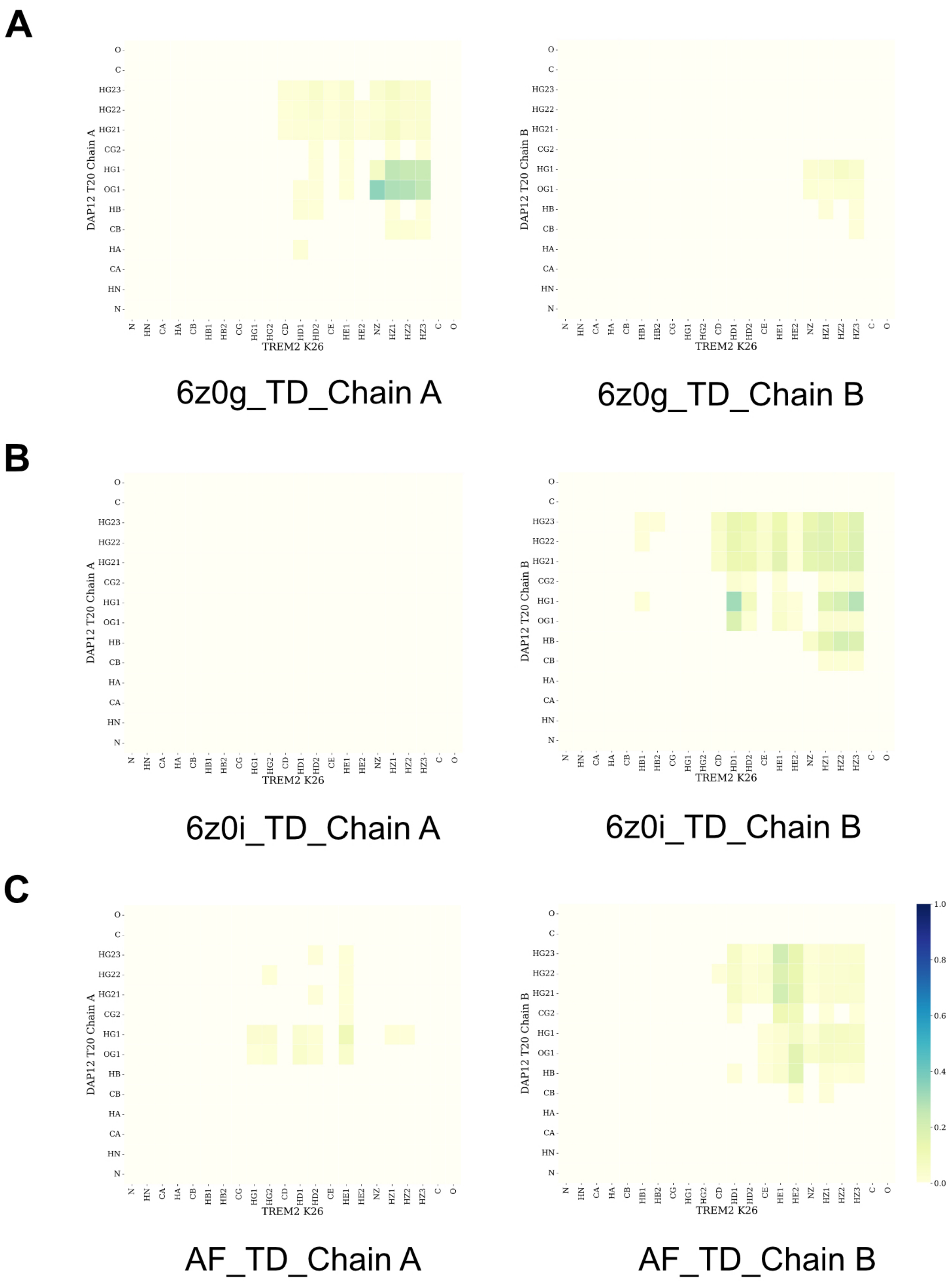


**Fig. S9.** Contact maps of K26/D16 residues.

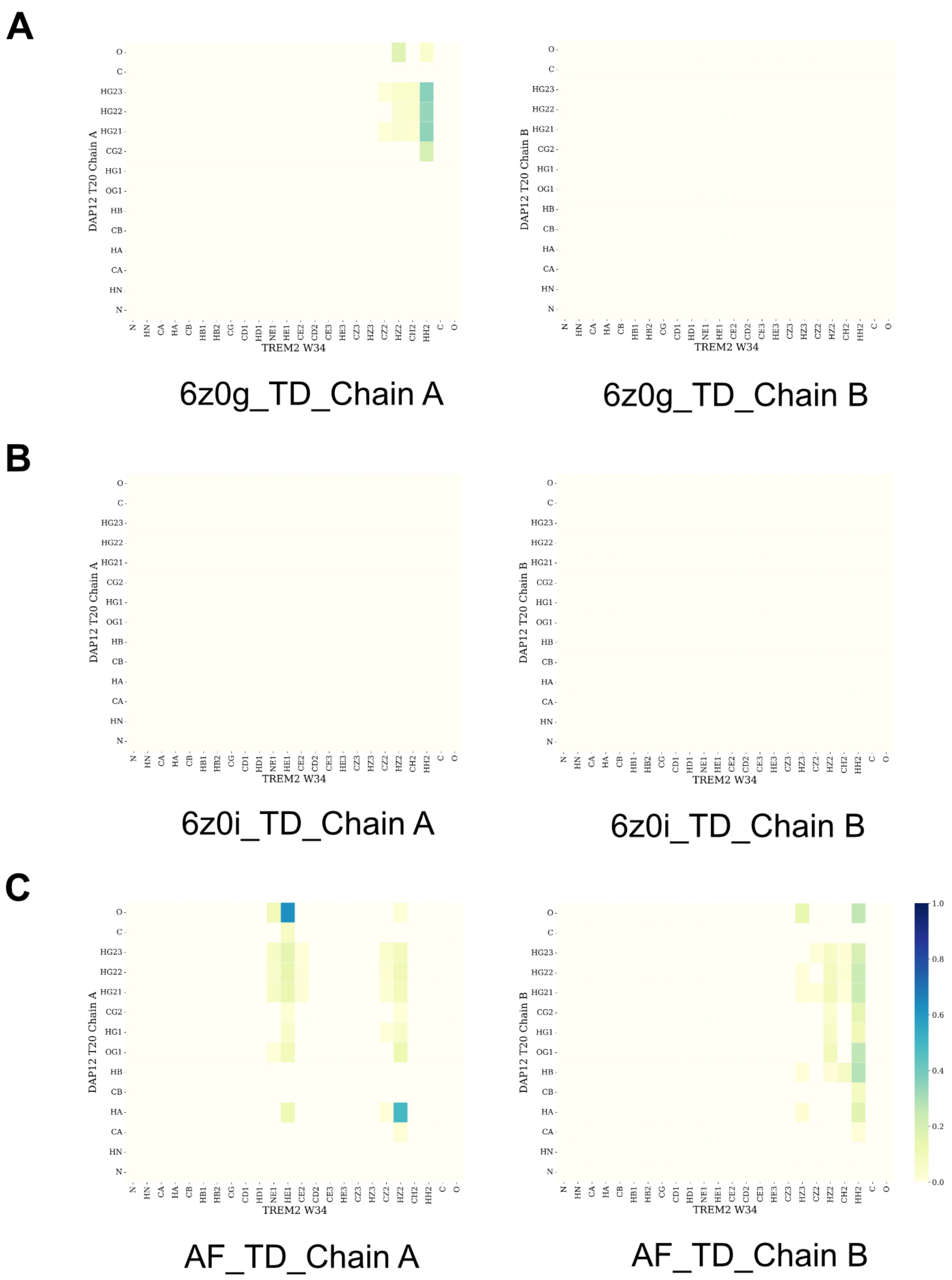


**Fig. S10.** Contact maps of K26/T20 residues.

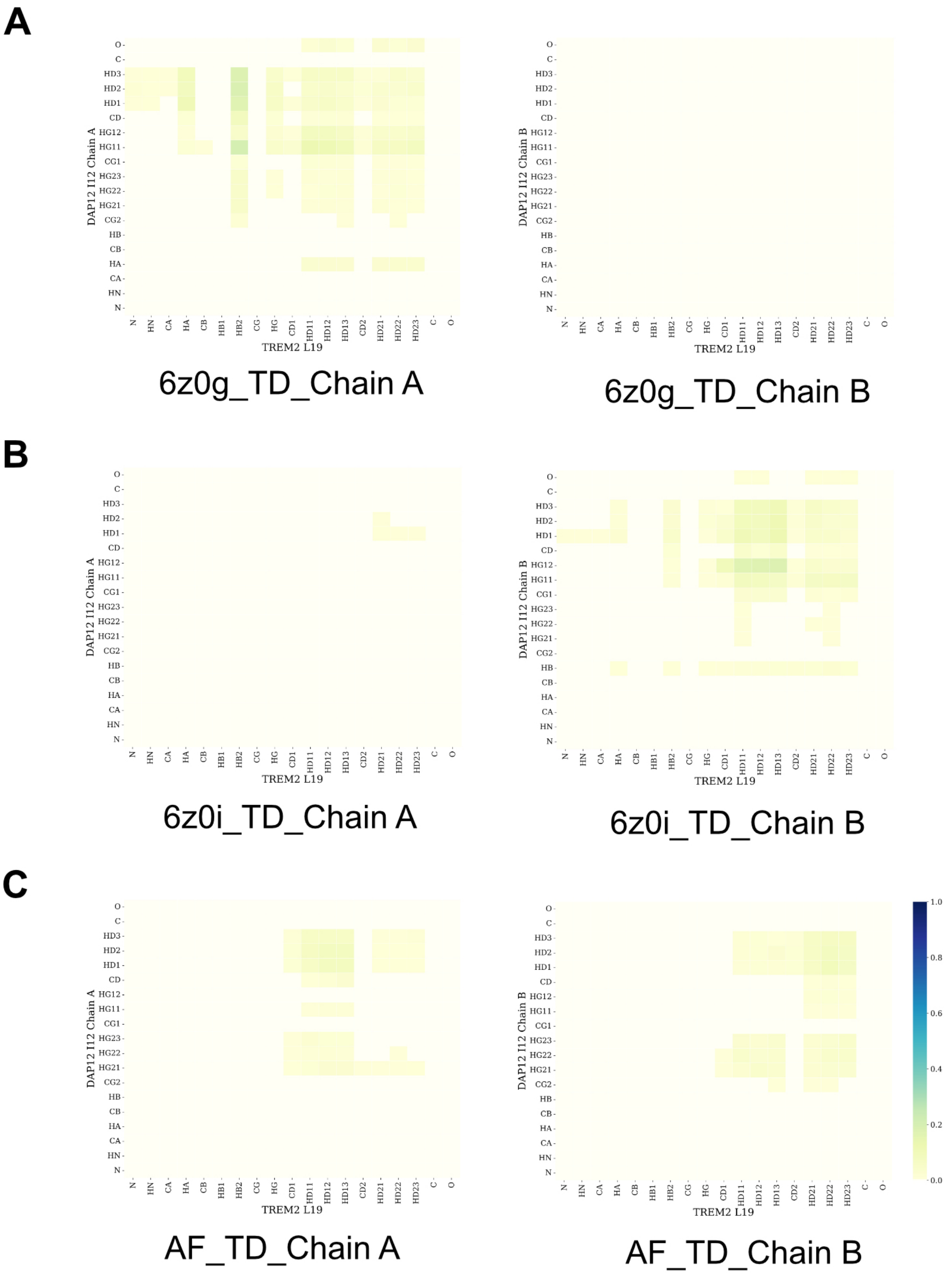


**Fig. S11.** Contact maps of W34/T20 residues.

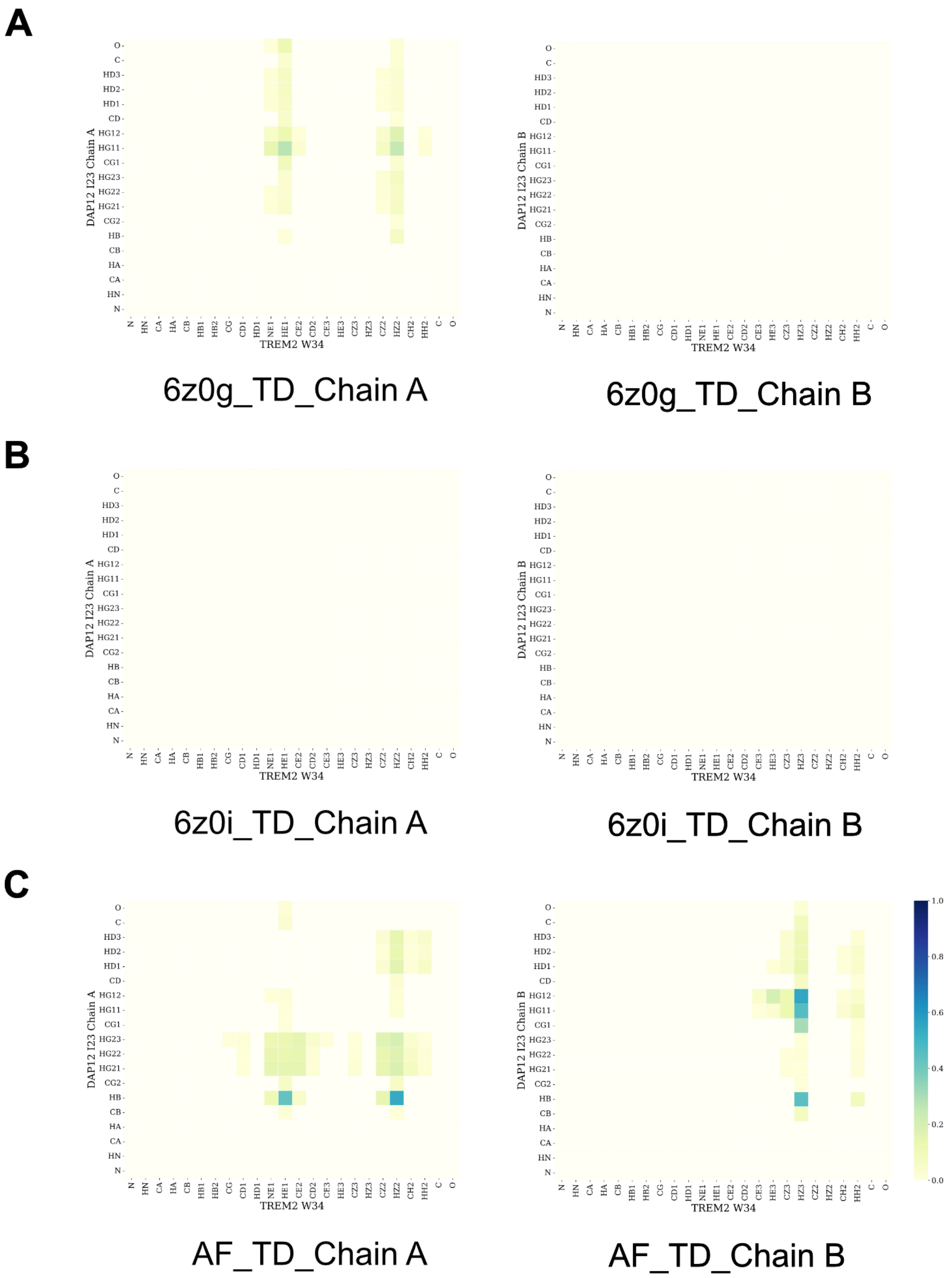


**Fig. S12.** Contact maps of W34/I23 residues.

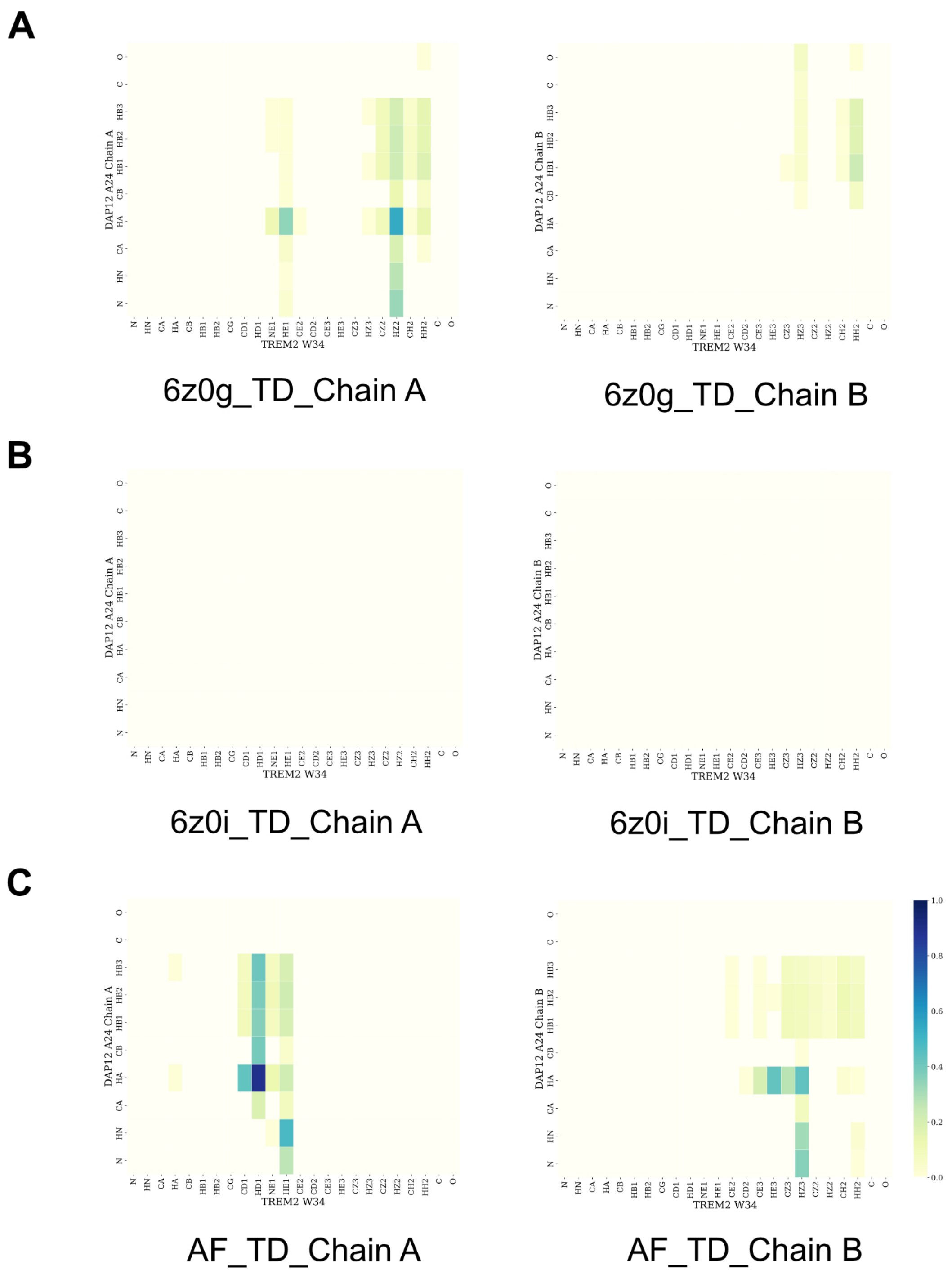


**Fig. S13.** Contact maps of W34/A24 residues.

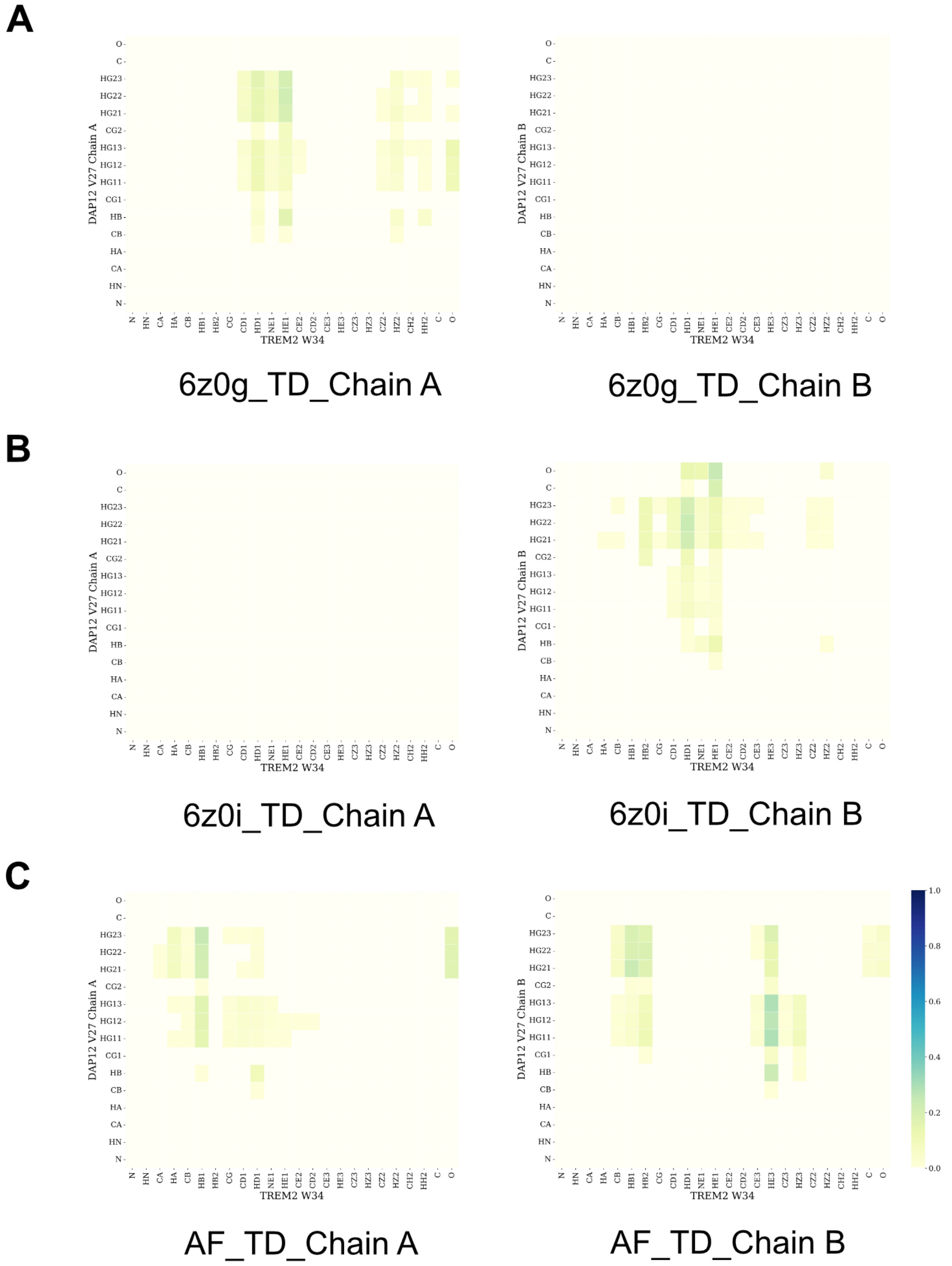


**Fig. S14.** Contact maps of W34/V27 residues.

 

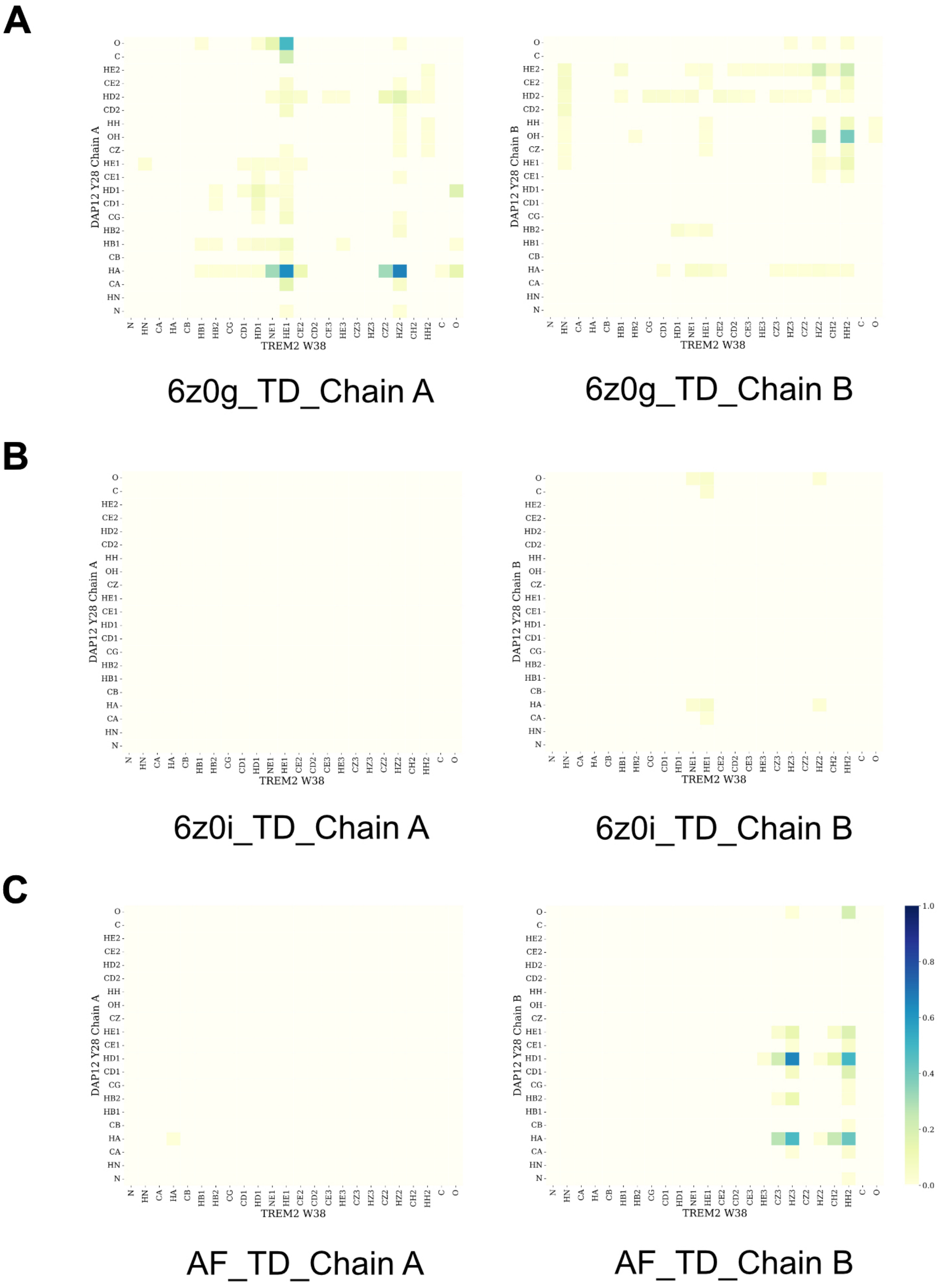


**Fig. S15.** Contact maps of W34/Y28 residues.

**Table S1. Hydrogen bonds possibility of K26/D16.**

| system | HZ1-A[a] | HZ2-A | HZ3-A | Total-A | HZ1-B[a] | HZ2-B | HZ3-B | Total-B |
|---|---|---|---|---|---|---|---|---|
| 6z0g_OD1 | 7.90 | 14.38 | 12.46 | 34.74 | 0 | 0 | 0 | 0 |
| 6z0g_OD2 | 20.20 | 9.52 | 14.85 | 44.57 | 0 | 0 | 0 | 0 |
| 6z0i_OD1 | 6.15 | 9.68 | 9.02 | 24.85 | 15.33 | 7.67 | 11.91 | 34.91 |
| 6z0i_OD2 | 14.77 | 16.90 | 28.68 | 60.35 | 19.83 | 11.23 | 11.97 | 43.03 |
| AF_OD1 | 10.23 | 10.30 | 12.33 | 32.86 | 8.77 | 12.67 | 11.85 | 33.29 |
| AF_OD2 | 13.33 | 11.89 | 13.27 | 38.49 | 11.95 | 12.85 | 9.98 | 34.78 |

[a]A is refered to Chain A. [b]B is refered to Chain B.

**Table S2. Hydrogen bonds possibility of K26/T20.**

| system | HZ1-A | HZ2-A | HZ3-A | Total-A | HZ1-B | HZ2-B | HZ3-B | Total-B |
|---|---|---|---|---|---|---|---|---|
| 6z0g_OG1 | 5.02 | 5.50 | 4.58 | 15.10 | 0.17 | 0.25 | 0.10 | 0.52 |
| 6z0i_OG1 | 0 | 0 | 0 | 0 | 0 | 0 | 0 | 0 |
| AF_OG1 | 0 | 0 | 0 | 0 | 0.35 | 0.35 | 0.32 | 1.02 |

**Table S3. Hydrogen bonds possibility of W34/T20.**

| system | HE1-A | HE1-B | system | HE1-A | HE1-B |
|---|---|---|---|---|---|
| 6z0g_OG1 | 0 | 0 | 6z0g_O | 0 | 0 |
| 6z0i_OG1 | 0 | 0 | 6z0i_O | 0 | 0 |
| AF_OG1 | 0.55 | 0 | AF_O | 9.08 | 0 |

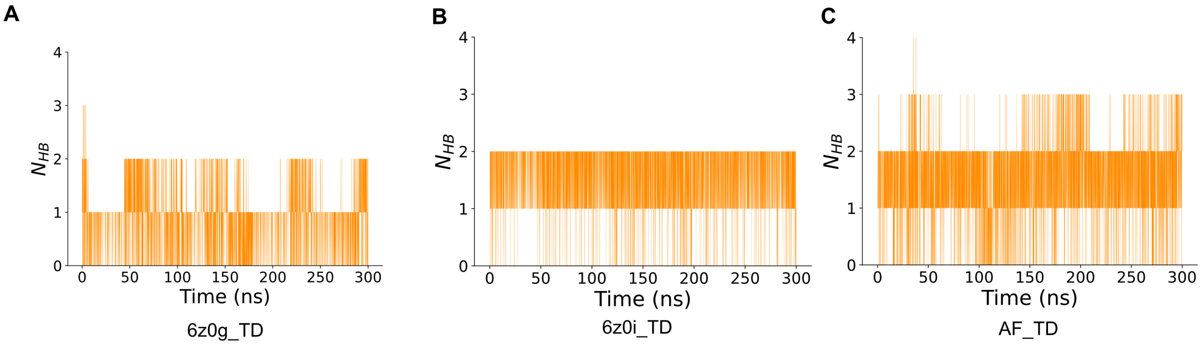


**Fig. S16.** Total number of hydrogen bonds of three systems.

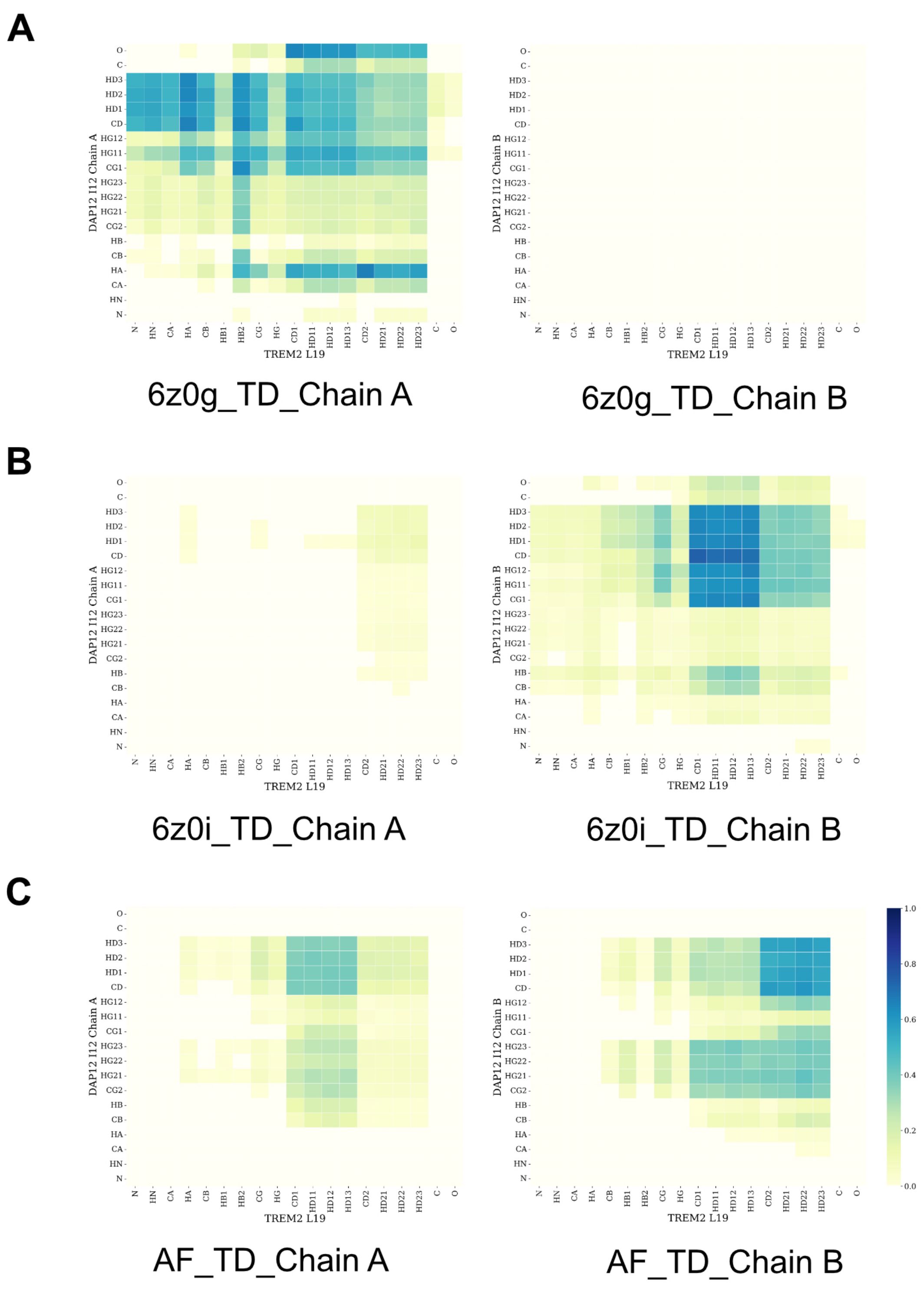


**Fig. S17.** Hydrophobic interaction L19/I12 in three systems.

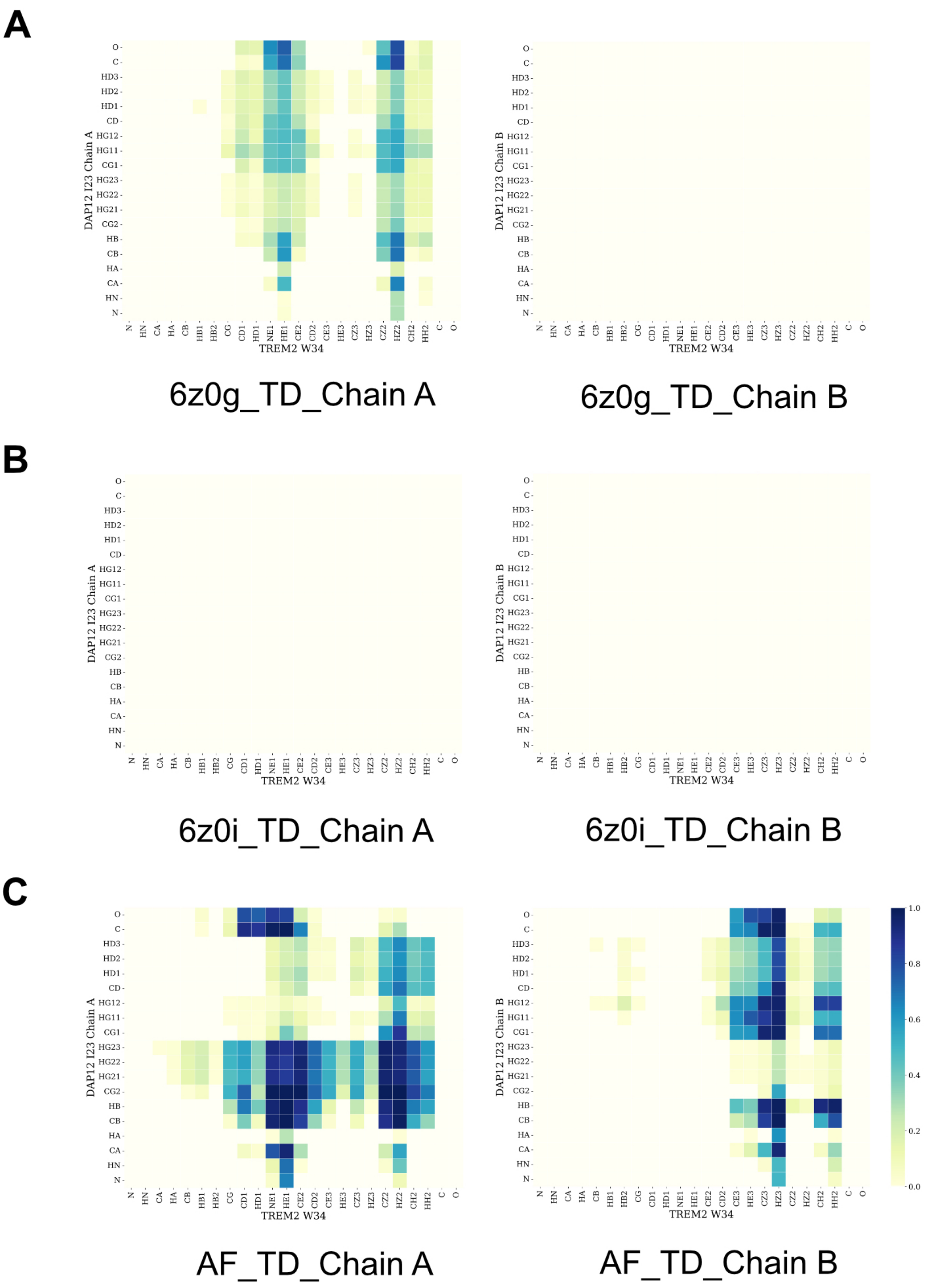


**Fig. S18.** Hydrophobic interaction W23/I23 in three systems.

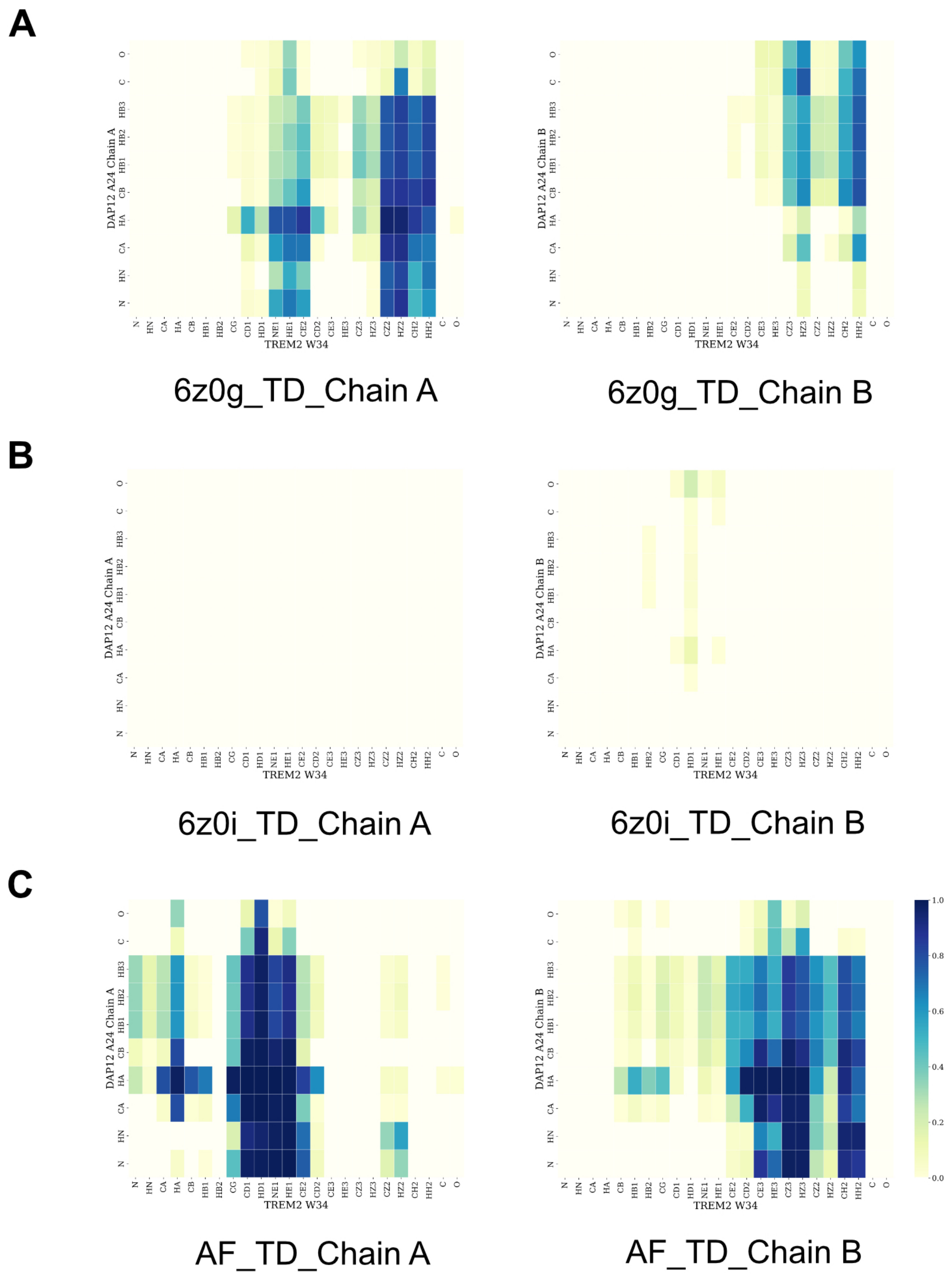


**Fig. S19.** Hydrophobic interaction W34/A24 in three systems.

 

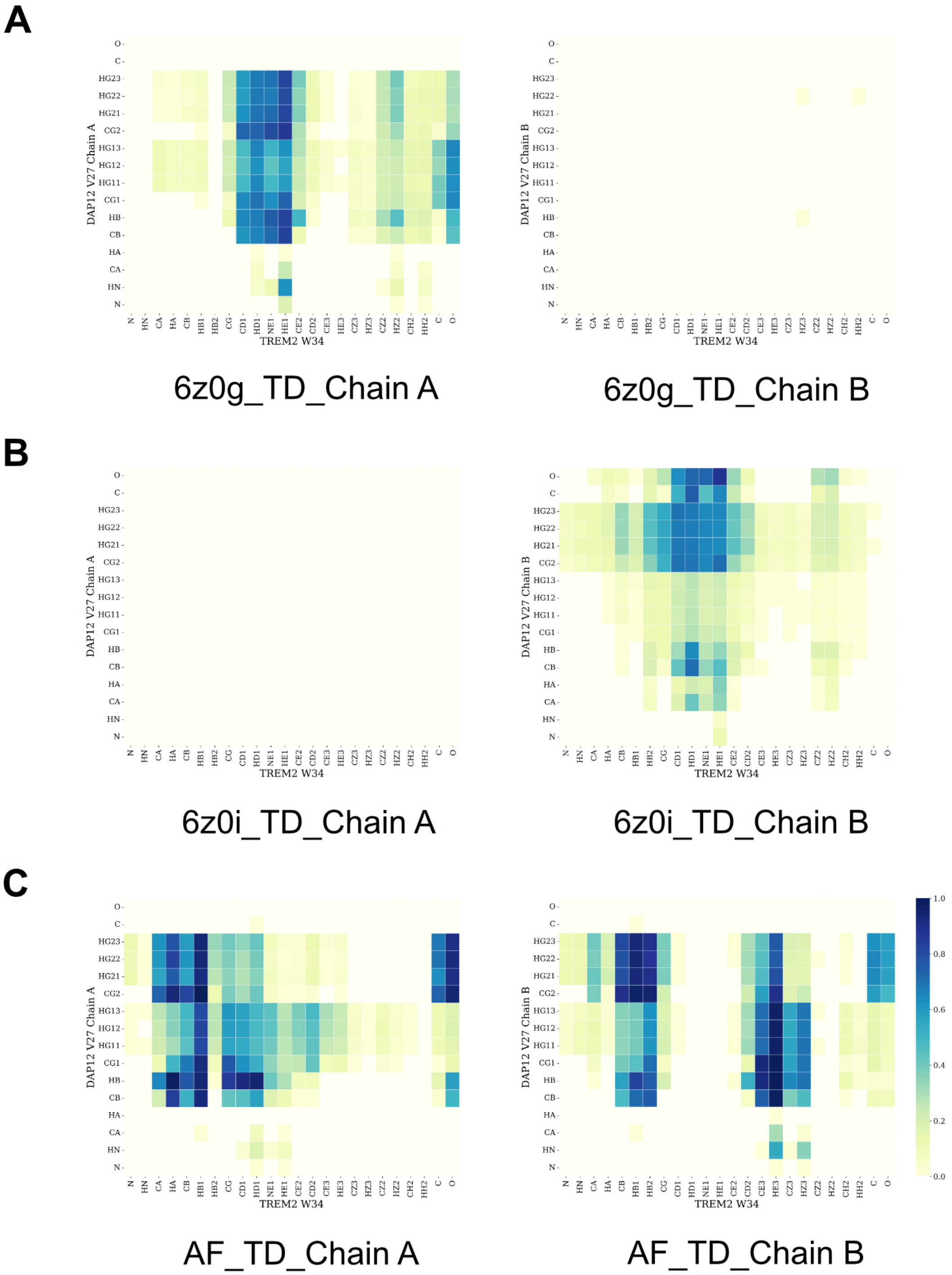


**Fig. S20.** Hydrophobic interaction W34/V27 in three systems.

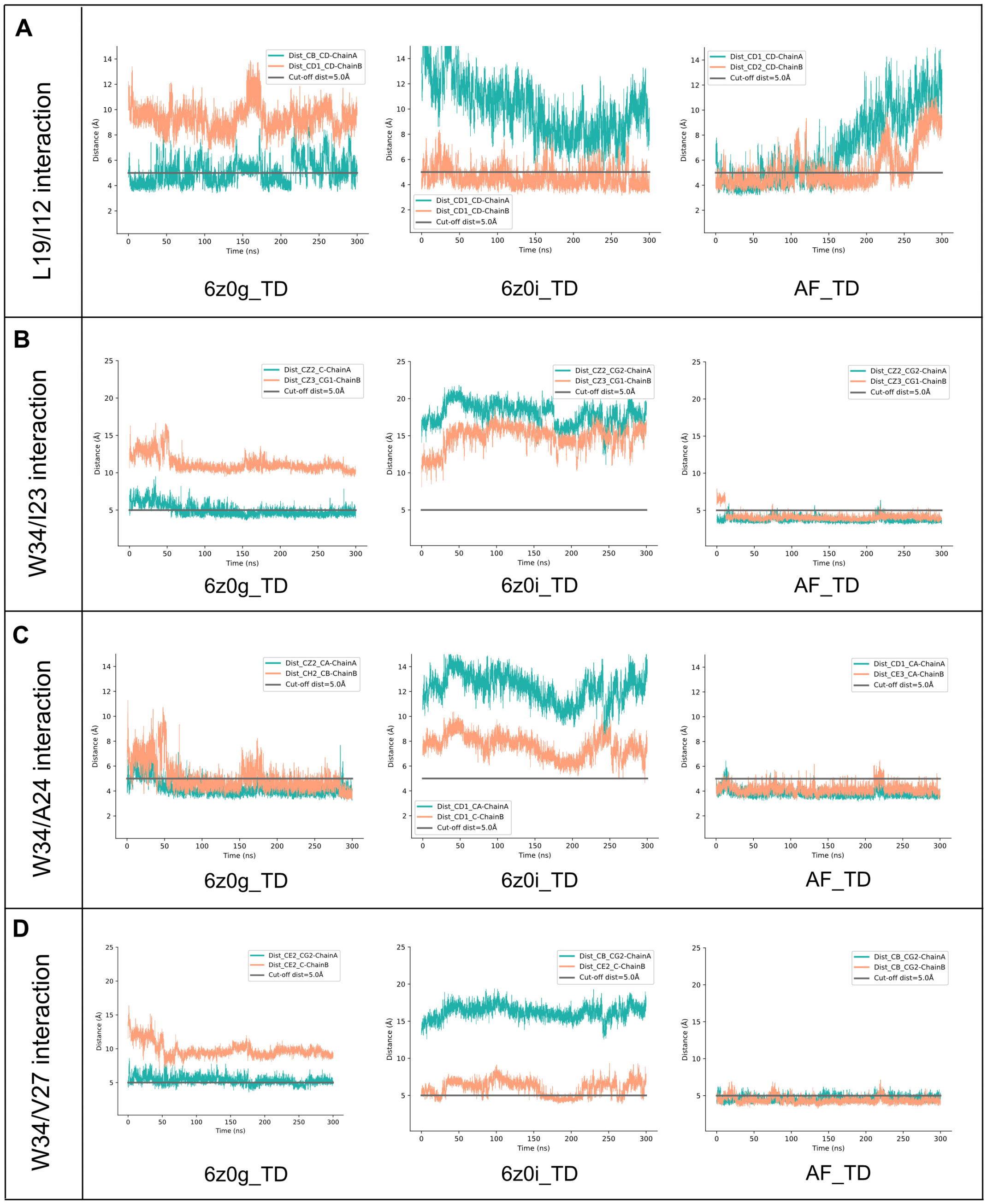


**Fig. S21.** Hydrophobic interaction of key residues in three systems. The cut-off distance is set as 11 Å.

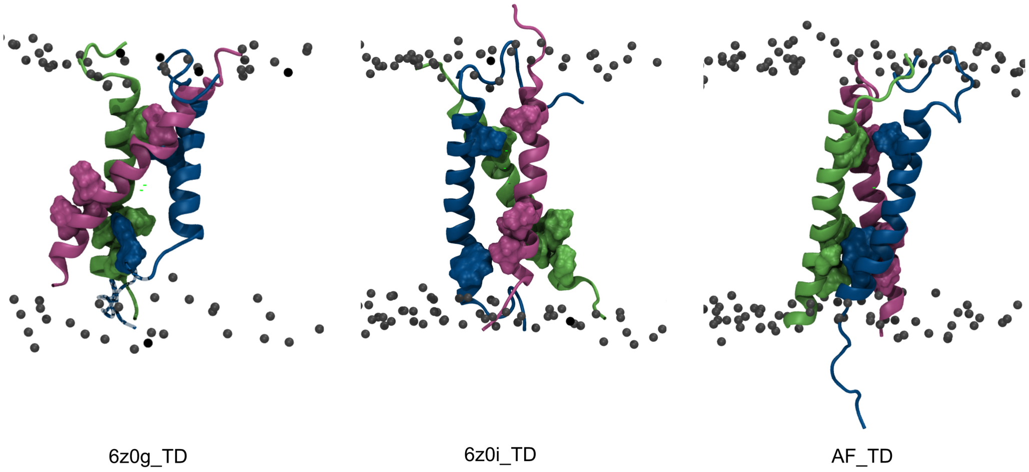


**Fig. S22.** Hydrophobic residues facing direction in three systems.

**Table S4. Sequence of three systems.**

| chains | sequence |
|---|---|
| DAP12 Chain A | *CSTVSPGVLAGIVVGDLVLTVLIALAVYFLGRL* |
| DAP12 Chain B | *CSTVSPGVLAGIVVGDLVLTVLIALAVYFLGRL* |
| TREM2 | *RSLLEGEIPFPPTSILLLLACIFLIKILAASALWAAAWHGQKPGTH* |
